# Prodrug activation in malaria parasites mediated by an imported erythrocyte esterase, acylpeptide hydrolase (APEH)

**DOI:** 10.1101/2024.08.30.610542

**Authors:** SA Sundararaman, JJ Miller, EC Daley, KA O’Brien, P Kasak, AM Daniels, RL Edwards, KM Heidel, DA Bague, MA Wilson, AJ Koelper, EC Kourtoglou, AD White, SA August, GA Apple, RW Rouamba, AJ Durand, JJ Esteb, FL Muller, RJ Johnson, GC Hoops, CS Dowd, AR Odom John

## Abstract

The continued emergence of antimalarial drug resistance highlights the need to develop new antimalarial therapies. Unfortunately, new drug development is often hampered by poor drug-like properties of lead compounds. Prodrugging temporarily masks undesirable compound features, improving bioavailability and target penetration. We have found that lipophilic diester prodrugs of phosphonic acid antibiotics, such as fosmidomycin, exhibit significantly higher antimalarial potency than their parent compounds (1). However, the activating enzymes for these prodrugs were unknown. Here, we show that an erythrocyte enzyme, acylpeptide hydrolase (APEH) is the major activating enzyme of multiple lipophilic ester prodrugs. Surprisingly, this enzyme is taken up by the malaria parasite, *Plasmodium falciparum*, where it localizes to the parasite cytoplasm and retains enzymatic activity. Using a novel fluorogenic ester library, we characterize the structure activity relationship of APEH, and compare it to that of *P. falciparum* esterases. We show that parasite-internalized APEH plays an important role in the activation of substrates with branching at the alpha carbon, in keeping with its exopeptidase activity. Our findings highlight a novel mechanism for antimicrobial prodrug activation, relying on a host-derived enzyme to yield activation at a microbial target. Mutations in prodrug activating enzymes are a common mechanism for antimicrobial drug resistance (2–4). Leveraging an internalized host enzyme would circumvent this, enabling the design of prodrugs with higher barriers to drug resistance.

**Significance:** Rising antimalarial drug resistance threatens current gains in malaria control. New antimalarial drugs are urgently needed. Unfortunately, many drug candidates have poor drug-like properties, such as poor absorbability in the gastrointestinal tract, or poor accumulation at the site of action. This can be overcome by prodrugging, the addition of prodrug groups which mask poor drug features until they are removed by an activating enzyme. Here, we show that a red blood cell enzyme, acylpeptide hydrolase, is taken up by malaria parasites and serves as the activating enzyme for multiple lipophilic ester prodrugs. Our findings highlight a novel mechanism for prodrug activation, which could be leveraged to design novel prodrugs with high barriers to drug resistance.

## Introduction

Malaria remains a significant threat to global public health, with hundreds of millions of cases and over half a million deaths occurring annually (5). The majority of cases and deaths occur in Africa and are caused by the parasite *Plasmodium falciparum*. Resistance to artemisinin-based therapies, current first line antimalarial regimens, has emerged in Southeast Asia and multiple parts of Africa (6–8). This emergence threatens current gains in global malaria control, and highlights the urgent need for new antimalarials with distinct mechanisms-of-action.

Phosphonic acid antibiotics, such as fosmidomycin (Fsm) (Fig. S1), represent a novel class of antimalarials. Fsm inhibits 1-deoxy-D-xylulose-5-phosphate reductoisomerase (DXR; E.C. 1.1.1.267), an enzyme in the apicoplast methylerythritol phosphate (MEP) pathway of isoprenoid biosynthesis (9). As the MEP pathway is found in apicomplexa, but absent in mammals, it represents an enticing target for drug development. However, while fosmidomycin is safe and well tolerated, it displays relatively poor antimalarial potency, a short serum half-life, and relatively poor bioavailability (1, 10). Although the antimalarial activity of fosmidomycin has been recognized since 1999, these shortcomings have contributed to suboptimal clinical efficacy and hindered development (11, 12).

We have previously found that lipophilic diester prodrugs of Fsm, and structurally related compounds, are highly potent antimalarials (1, 13). One such prodrug, POM-ERJ (RCB-185), which contains a lipophilic pivaloyloxymethyl (POM) promoiety, exhibited 10-fold higher potency than its parent compound (the Fsm analog ERJ) (Fig. S1) both *in vitro* and in a mouse model of malaria (1). We hypothesized that the increased potency of POM-ERJ in cell culture is due to improved cellular penetration, increasing drug concentrations in the parasite. Prodrugs are inactive against their target enzyme, and are typically converted to the active drug by enzymatic removal of the promoiety. If the potency of POM-ERJ is correlated with its ability to enter the parasite, this implies that POM-ERJ is de-esterified and activated by an intraparasitic enzyme. Identification of the enzyme responsible for POM-ERJ activation could aid in the design of novel prodrugs that are more rapidly or specifically activated by that enzyme, further increasing their antimalarial activity or selectivity.

In this study, we show that an erythrocyte enzyme, acylpeptide hydrolase (APEH), is internalized by *P. falciparum* and responsible for the majority of activation of the antimalarial prodrug POM-ERJ. Our findings provide insight into prodrug activation within the parasite, and raise the possibility of designing host enzyme-activated prodrugs with a high barrier to drug resistance.

## Results

### Erythrocyte acylpeptide hydrolase (APEH) displays esterase activity within the *P. falciparum* cytoplasm

We previously used drug resistance screens to identify a POM-ERJ-activating enzyme (hydroxyacylglutathione hydrolase, GloB) in zoonotic staphylococci and *Staphylococcus aureus* (14). We sought to identify a POM-ERJ-activating enzyme in *P. falciparum* malaria parasites using a similar forward chemical genetics approach (14, 15). We exposed wild type strains, as well as strains with partial resistance to ERJ and Fsm, to varying doses of POM-ERJ (0.25-5x EC_50_), but failed to generate resistant lines (16). We attempted an intermittent drug pressure approach, again with negative results, in contrast to the ease with which we have previously been able to generate fosmidomycin-resistant lines (16, 17). We therefore sought an alternative approach to identify putative prodrug-activating enzymes in malaria parasites.

Lipophilic esters are typically hydrolyzed by serine hydrolases, a superfamily of enzymes that includes esterases, proteases, peptidases, and lipases (18). The *P. falciparum* genome contains at least 43 annotated serine hydrolases, of which 22 are predicted to be essential based on knockout screens (19–22). As generating individual knockout or knockdown lines of each serine hydrolase was infeasible, we leveraged a 96-compound library of fluorescent ester substrates to identify candidate prodrug activating enzymes based on the enzymes’ biochemical activity (23). This library consists of fluorescein derivatives wherein a reactive ester moiety has been added (Fig. 1A, Fig. S2) (24). These esters mask fluorescence until they are removed by hydrolysis. Thus, substrate de-esterification is proportional to the increase in fluorescence over time.

**Figure 1.**
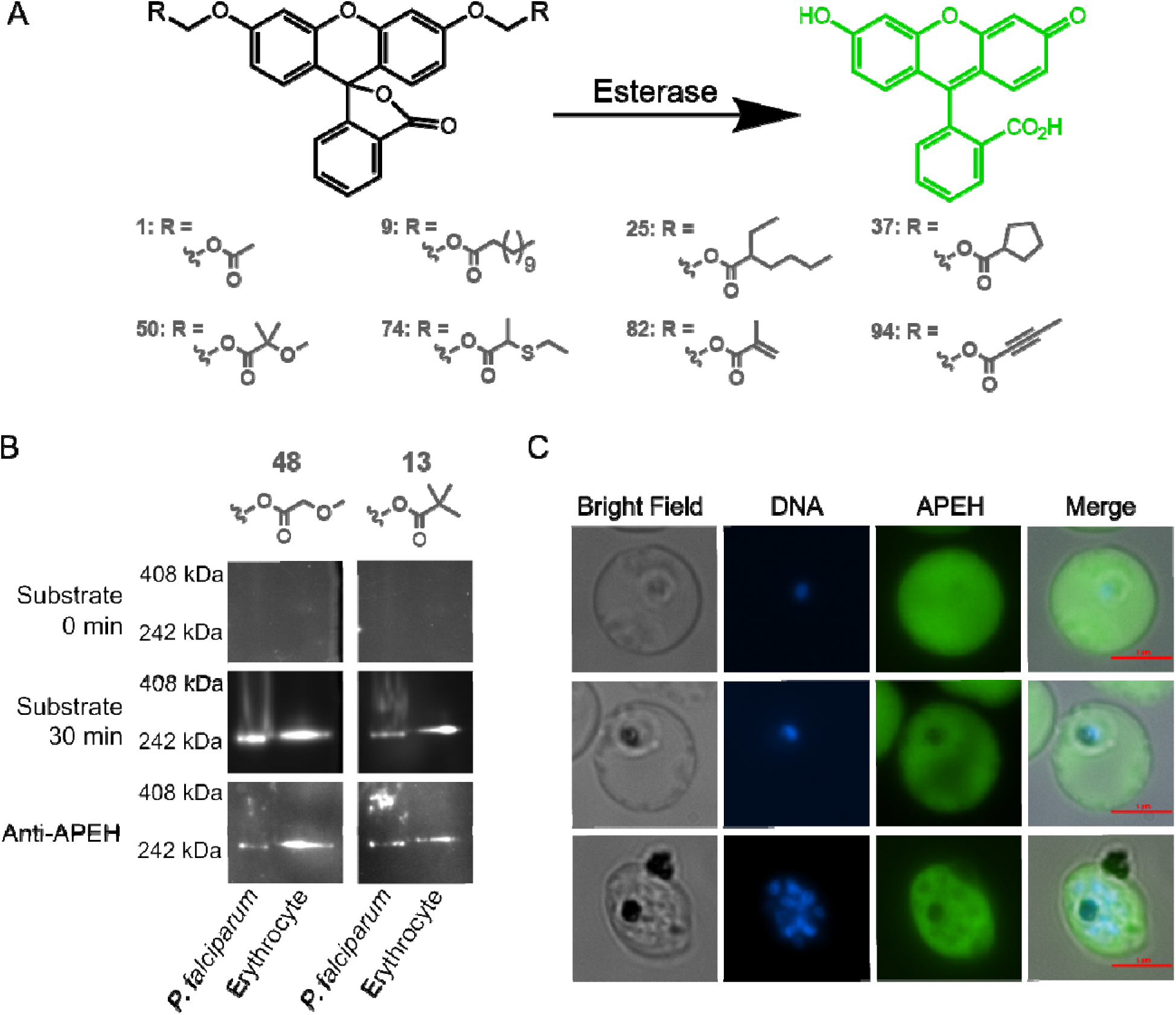
Human APEH is active in *P. falciparum* lysate and localizes to the cytoplasm of malaria parasites. **A**) Schematic of the fluorescent substrate library used in this study. Reactive ester moieties (R) are added to fluorescein yielding substrates with masked fluorescence. Hydrolysis of the ester bond by an esterase produces a fluorescent signal. A subset of substrates, demonstrating the various substrate types found in the library, is shown here. The full 96 substrate library is shown in Fig. S2. **B**) Native in gel fluorescence and immunoblotting of *P. falciparum* or erythrocyte lysate using substrates 48 and 13 reveals a single prominent band of esterase activity corresponding to the human enzyme, APEH. Gels are shown prior to incubation with fluorescent substrate (0 min) and after 30 minutes of incubation. Proteins from each gel were then transferred nitrocellulose membranes for immunoblotting with anti-APEH. **C**) Immunofluorescence images of infected erythrocytes showing the cytoplasmic localization of APEH.

We searched for candidate esterases using native in-gel fluorescence. We separated lysate from intraerythrocytic stage *P. falciparum* and uninfected erythrocytes by native polyacrylamide gel electrophoresis (PAGE), and incubated gels with two fluorescent substrates—a promiscuous alkyl ether substrate (**48**), and a pivaloyloxymethyl (POM)-containing substrate (**13**) (Fig. 1B, Fig. S2) (15, 23). Both substrates produced a single dominant band of esterase activity, migrating at approximately 250 kDa. Surprisingly, this single dominant band was present in both *P. falciparum* lysate and lysate from uninfected erythrocytes (Fig. 1B, Fig. S3A). Analysis of the parasite lysate band by peptide mass-spectroscopy identified 127 *P. falciparum* proteins, none of which were predicted to be serine hydrolases (Table S1). Given the presence of a band with similar electrophoretic mobility in erythrocyte lysate, we also searched for matches to human proteins. We identified 124 human proteins, including one serine hydrolase, acylpeptide hydrolase (APEH) (Table S1). We confirmed the presence of APEH at the site of fluorescence by immunoblotting (Fig. 1B, Fig. S3A).

APEH is a ubiquitously expressed serine protease (25). The enzyme is thought to play a critical role in protein degradation, by removing terminal amino acids from damaged proteins (26). Human APEH has a molecular weight of 81 kDa, and forms a homo-tetramer, consistent with the migration pattern observed on native PAGE (27). Interestingly, recent studies have shown that APEH is internalized by *P. falciparum* and may be essential to asexual parasite growth (22, 28).

Prior studies of internalized APEH were unable to determine its subcellular localization within the parasite. Using indirect immunofluorescence, we found that APEH is localized to the parasite cytoplasm (Fig. 1C). The fluorescence intensity within the parasite was similar to that within the erythrocyte cytoplasm, and did not appear to vary throughout the asexual lifecycle. However, APEH was not detected in the food vacuole or the parasite nucleus. We independently confirmed this localization pattern using a second antibody to APEH (Fig. S3B). These data suggest that APEH accumulates in the *P. falciparum* cytoplasm, starting early after invasion of the erythrocyte and remains present even after schizogony (nuclear division).

### APEH activates POM-ERJ *in vitro*

As APEH hydrolyzes ester substrates, including pivaloyloxymethyl, we queried whether it could also de-esterify and activate the antimalarial POM-ERJ. We examined POM-ERJ activation using a target enzyme (DXR) activity assay. As prodrugs such as POM-ERJ must be activated through removal of the prodrug moiety to inhibit their target, this assay follows decreases in DXR activity to quantify prodrug activation (Fig. 2A, Fig. S4). Incubation of POM-ERJ with *P. falciparum* or uninfected erythrocyte lysate resulted in complete DXR inhibition, suggesting that both parasites and erythrocytes contain one or more enzymes that can activate POM-ERJ (Fig. 2B). Similarly, incubation of POM-ERJ with purified recombinant APEH (rAPEH) also led to complete DXR inhibition, providing strong support that APEH alone was sufficient for full prodrug activation. To determine if POM-ERJ-activating activity in parasite and erythrocyte lysate was attributable to APEH, we incubated lysate or rAPEH with a well characterized, irreversible, APEH inhibitor, AA74-1 (Fig. S1) (29). Addition of AA74-1 to rAPEH, parasite lysate, or erythrocyte lysate completely rescued DXR activity (Fig 2B). Together these data suggest that APEH activity is required for POM-ERJ activation in *P. falciparum* and erythrocytes (29).

**Figure 2.**
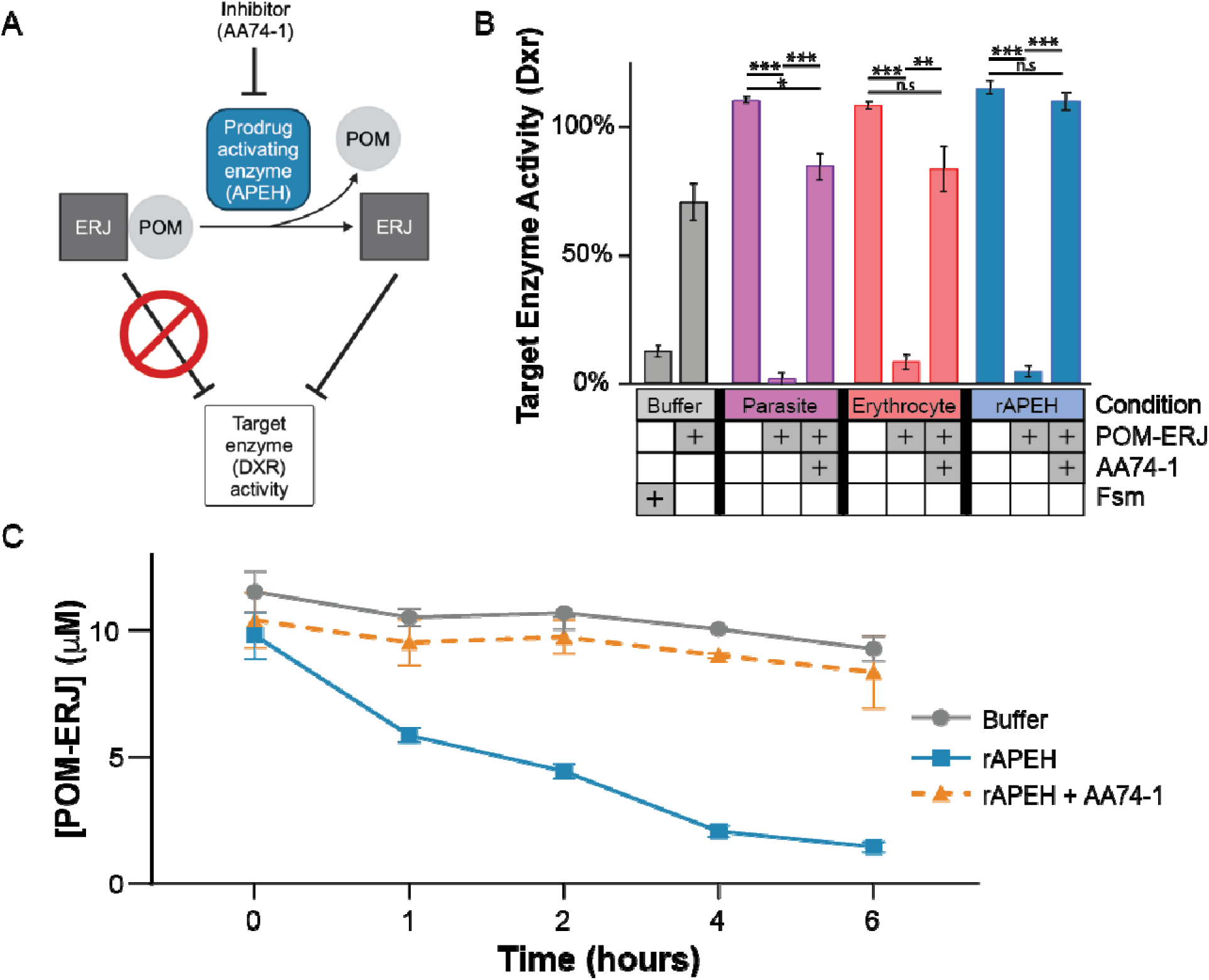
The human erythrocyte esterase APEH de-esterifies and activates POM-ERJ. **A**) Schematic of linked enzyme activity assay. Intact POM-ERJ does not inhibit DXR. Hydrolysis of POM-ERJ by a prodrug activating enzyme (APEH) yields active ERJ inhibitor, leading to inhibition of DXR. Inhibition of the APEH by its own inhibitor (AA74-1) impedes POM-ERJ hydrolysis, rescuing DXR activity. **B**) POM-ERJ activation as measured by inhibition of DXR activity. Parasite lysate, erythrocyte lysate, recombinant APEH (rAPEH), or buffer were incubated for 12 hours at 37°C in the presence or absence of POM-ERJ and AA74-1, and then added to recombinant DXR. Residual DXR activity is shown for each condition. Fosmidomycin (Fsm) is included as a positive control for DXR inhibition. Slight DXR inhibition is observed when POM-ERJ is incubated with buffer, likely due to slow spontaneous hydrolysis. **C**) Change in POM-ERJ concentration upon incubation with rAPEH, rAPEH and 400 nM AA74-1, or buffer, as measured by LC-MS. Data in B and C represent the mean of three replicates, error bars show SEM. P-values were calculated using Welch’s t-test. *\*\*\*: p<0.001, **: p<0.01, *: p<0.05, n.s. not significant*.

We next used liquid chromatography-mass spectrometry (LC-MS) to quantify the rate of POM-ERJ hydrolysis upon incubation with rAPEH. Incubation of 10 µM POM-ERJ with rAPEH resulted in near complete prodrug hydrolysis within 6h (Fig. 2C). Minimal hydrolysis was observed when POM-ERJ was incubated with buffer alone, or with rAPEH inhibited by AA74-1, further confirming the ability of APEH to rapidly activate POM-ERJ *in vitro*.

### Inhibition of APEH decreases POM-ERJ activity against *P. falciparum*

Having shown that APEH activates POM-ERJ *in vitro*, we wanted to determine if APEH functions as a prodrug-activating enzyme in a cellular context. As the APEH inhibitor AA74-1 also inhibits *P. falciparum* growth with nanomolar potency, we initially attempted to measure synergy or antagonism between AA74-1 and POM-ERJ using isobologram analysis, but were unable to detect a consistent effect (28). As AA74-1 irreversibly inhibits APEH, we tested if pre-incubation of erythrocytes with AA74-1 could produce a more reproducible effect. While pretreatment of erythrocytes followed by washout of unbound AA74-1 led to near complete inhibition of APEH, the anti-parasitic effect was substantially ameliorated (Fig. S5). This allowed us to specifically measure the effects of APEH inhibition on POM-ERJ activation in living cultured malaria parasites.

To test the necessity of APEH for activation of POM-ERJ in a cellular context, we continuously cultured *P. falciparum* in parasites that had been pretreated with 1 uM AA74-1, yielding near total inhibition of APEH (APEH-inhibited, Fig. S5B), or DMSO (APEH-active). We then performed dose-response analyses of POM-ERJ in APEH-inhibited and APEH-active cultures (Fig. 3). Parasites grown in APEH-inhibited erythrocytes displayed increased resistance to POM-ERJ, as demonstrated by a 2.7-fold increase in the half-maximal effective concentration (EC_50_) (Fig. 3). APEH inhibition had no effect on the EC_50_ of fosmidomycin, suggesting that the effect on POM-ERJ was due to inhibition of prodrug activation rather than another mechanism (Fig. 3). These data suggest that APEH is the predominant POM-ERJ-activating enzyme in *P. falciparum* culture.

**Figure 3.**
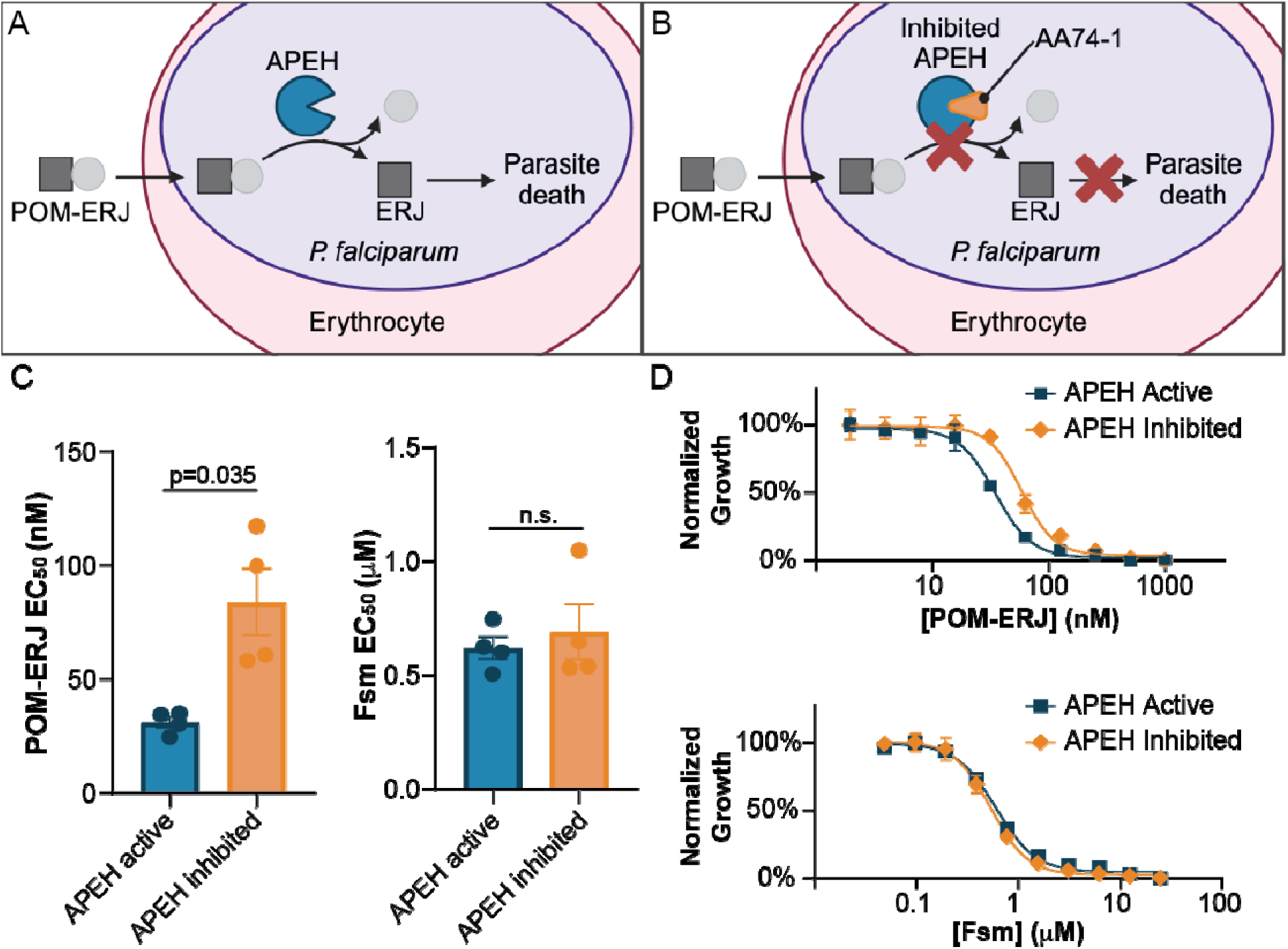
Erythrocyte APEH is required for *P. falciparum* sensitivity to POM-ERJ. (**A, B**) Effect of APEH inhibition on POM-ERJ activation. Active APEH (**A**) hydrolyzes POM-ERJ, leading to accumulation of active drug (ERJ) in *P. falciparum* and parasite death. APEH inhibition by AA74-1 (**B**) decreases POM-ERJ activation, decreasing ERJ accumulation and parasite death. **C**) EC_50_ values for POM-ERJ or Fsm against *P. falciparum* 3D7 grown in AA74-1 treated (APEH inhibited) or DMSO treated (APEH active) erythrocytes. Bar graphs (mean ± SEM) are representative of four biological replicates. Points represent EC_50_s for each replicate. For POM-ERJ, APEH active cultures have an EC_50_ of 31.0 ± 2.4 nM and APEH inhibited cultures an EC_50_ of 83.9 ± 14.7 nM. For Fsm, APEH active cultures have an EC_50_ of 0.62 ± 0.04 μM and APEH inhibited an EC_50_ of 0.69 ± 0.12 μM. P-values were calculated using Welch’s t-test. No statistical significance is denoted by n.s. (**D**) Representative dose-response curves for POM-ERJ and Fsm against *P. falciparum* 3D7. Data are representative of two technical replicates. Points represent mean growth ± SEM at each concentration.

### APEH structure activity relationship and activation of a distinct POM ester prodrug

We recently found that a structurally distinct POM prodrug (POM-HEX) inhibits the glycolytic enzyme enolase in erythrocytes and displays antimalarial activity (30). Given the shared prodrug moiety, we wondered if APEH could also activate POM-HEX. We measured POM-HEX activation by monitoring the rate of glycolysis in erythrocytes using the Agilent Seahorse XFe96 Pro Analyzer. This instrument estimates rates of glycolysis in live cells by measuring the extracellular acidification rate in a transient microchamber environment (Fig. S6A). Treatment of erythrocytes with POM-HEX resulted in a gradual decrease in glycolysis relative to controls (Fig. S6B). This decrease was not seen when erythrocytes were pretreated with AA74-1, suggesting that APEH activates POM-HEX in erythrocytes.

To more fully evaluate the structure-activity relationship of APEH, and compare it to that of other *P. falciparum* serine hydrolases, we quantified the rates of activation of additional substrates from our fluorescein-based ester substrate library (23, 31). Rates of substrate activation by APEH and parasite lysate were positively correlated (Fig. 4, Table S2). A notable exception were long chain unbranched and branched alkyl esters, where activation by APEH was not detectable in our assay. Similarities between rates of substrate activation by parasite lysate and APEH could indicate either that APEH is the major activating enzyme for a given substrate, or that the substrate is activated by multiple enzymes (including APEH). To more clearly identify which substrates were predominantly activated by APEH present in parasite lysate, we measured substrate activation after treatment of parasite lysate with AA74-1 (Fig. 4, Table S2). Substrates with a cyclic group or branching at the alpha carbon showed the lowest residual activity after AA74-1 treatment, suggesting that APEH is the major activating enzyme for these substrates in parasite lysate.

**Figure 4.**
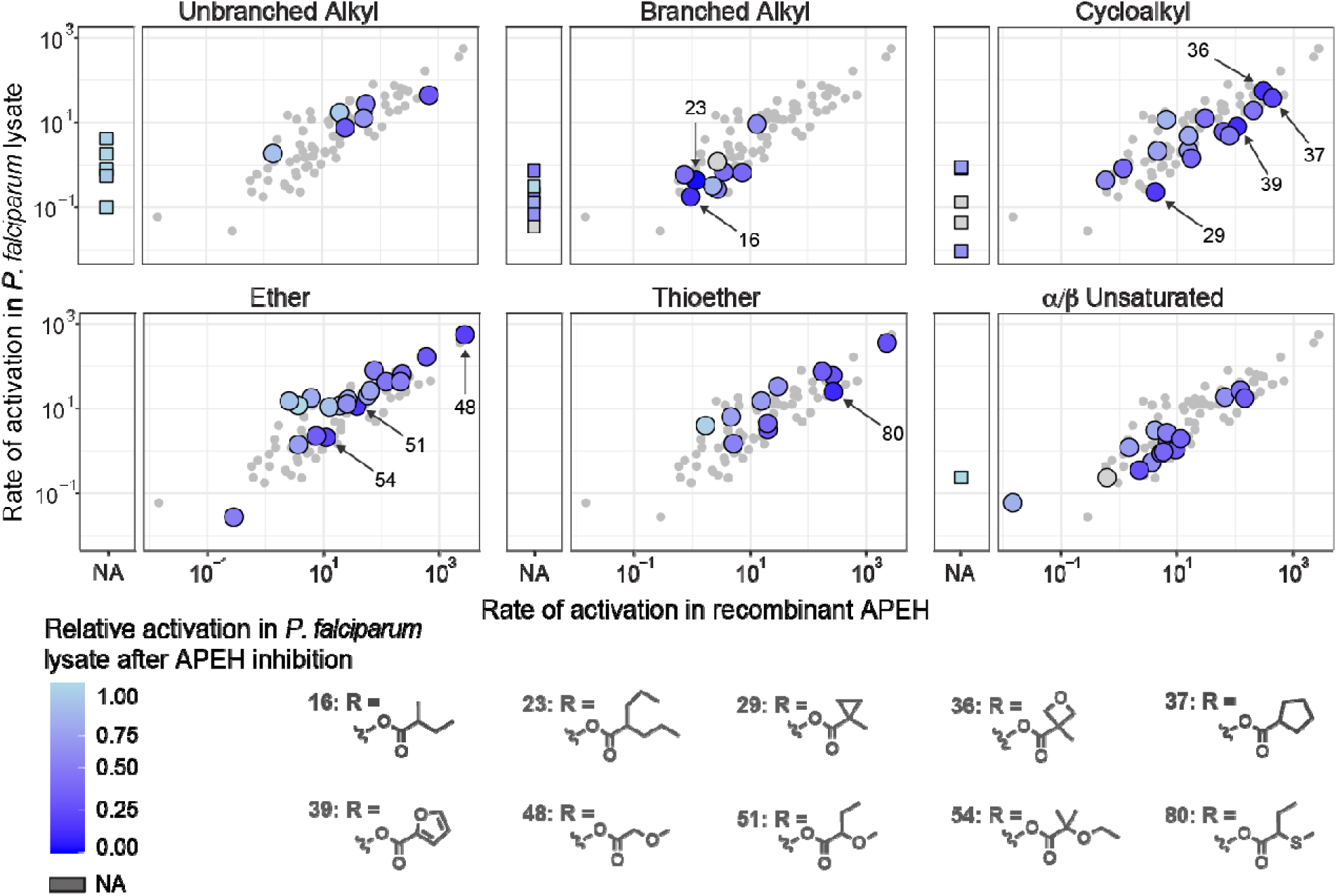
Structure activity relationship of recombinant APEH and *P. falciparum* lysate with and without APEH inhibition. Rates of fluorescent substrate activation (fM fluorescein produced per minute per ng protein) are shown for purified recombinant APEH (x-axis) and soluble parasite lysate (y-axis). Substrates are grouped by promoiety type, with large circles representing substrates within that type, and small circles representing the full library for comparison. Substrates with no detectable activation by APEH are shown as squares to the left of each main plot. Colors represent the relative rate of substrate activation in *P. falciparum* lysate after AA74-1 treatment (white: no change, dark blue: complete inhibition, dark gray: no data). The ten substrates with the highest decrease in activation after APEH inhibition are shown (activation rates for all individual substrates are presented in Table S2).

## Discussion

Many FDA-approved prodrugs in clinical use today are thought to be activated by enzymes in the host liver or serum. In recent years, prodrugs that are activated by tissue- or pathogen-specific enzymes have gained increased interest (2, 32–36). Designing pathogen-specific prodrugs would allow these compounds to maintain beneficial prodrug properties (*i.e.*, membrane permeability) until reaching their target, thus increasing drug potency while decreasing off-target effects. Here, we present an unexpected finding: that a host enzyme, APEH, localized within the *P. falciparum* cytoplasm, remains catalytically active and serves as a prodrug-activating enzyme. Our data show that APEH is critical to POM-ERJ activation in *P. falciparum* culture, with APEH inhibition significantly reducing the sensitivity of *P. falciparum* to POM-ERJ.

While the potency of POM-ERJ is reduced by APEH inhibition, the prodrug continues to inhibit growth at nanomolar potency. This could be due to low level residual APEH activity, which was detectable at 48 and 72 hours after washout (Fig. S5B), as well as slow spontaneous hydrolysis. It is also possible that other esterases can activate POM-ERJ. We note that activation of fluorogenic substrate **13**, which contains a POM progroup, was significantly but incompletely inhibited by AA74-1 (∼50% inhibition, Fig. 4, Table S2). Together, these findings suggest that, while APEH is the predominant activating enzyme of this substrate, other *P. falciparum* esterases may also have the capacity to activate POM prodrugs.

Leveraging an internalized host enzyme for pathogen-targeted prodrug activation presents both advantages and disadvantages. Prodrugs activated by pathogen-encoded enzymes risk drug resistance due to selection for mutations in the activating enzyme (Fig. S7). For example, mutations in the *P. falciparum* enzyme *PfPARE* have been shown to increase resistance to multiple candidate antimalarials by disrupting prodrug activation (2–4). Similarly, we recently found that mutations in the staphylococcal esterases GloB and FrmB can lead to antibacterial resistance to POM prodrugs (14, 15). Activation by a host-encoded enzyme would circumvent this risk, as is highlighted by our inability to generate *P. falciparum* lines that are resistant to POM-ERJ. At the same time, activation by a host enzyme could reduce prodrug half-life and increase off-target effects, due to activation in the blood or other tissues. Prior work in our lab found that POM-ERJ was effective in mouse models of malaria, suggesting that benefits of increased membrane permeability may outweigh decreases in serum half-life (1). Moreover, as phosphonate antimalarials target an enzyme with no human homolog, activation at other sites would not necessarily increase adverse effects.

Using a large fluorogenic substrate library, we identify a number of ester substrates that are readily hydrolyzed by APEH. Our data show that inhibition of APEH in parasite lysate significantly decreases activation of substrates with branching at the alpha carbon. We suspect that this is related to the exopeptidase activity of APEH, as the side chains of amino acids branch from the alpha carbon. These findings could serve as the basis for the development of new prodrugs with increased activation by APEH, or, alternatively, that resist hydrolysis by APEH.

Importantly, while we examined the activity of full length recombinant APEH, evidence suggests that the enzyme is proteolytically cleaved during or after import into the parasite (28).

Supporting this, we noted that parasite-localized APEH migrates slightly faster than erythrocyte localized-APEH in native PAGE. This change in electrophoretic mobility could indicate that cleavage results in a conformational change or loss of a portion of the enzyme. Prior studies of APEH in the bovine and human eye have shown that enzymatic cleavage by trypsin alters the substrate preference of APEH. If this is also the case for parasite-internalized APEH, the differential substrate activity of cleaved and intact enzyme could lead to the development of prodrugs that are specifically activated by the internalized form of the enzyme.

Our data suggest that both erythrocyte-localized and parasite-internalized APEH de-esterify and activate POM-ERJ. As the membrane permeability of POM-ERJ likely contributes to its increased potency against *P. falciparum*, we suspect that the majority of POM-ERJ activation occurs within the parasite itself, although this could not be experimentally confirmed. It is also possible that POM-ERJ is activated within the erythrocyte, and active ERJ then enters the parasite by another mechanism. Further elucidation of sites of prodrug activation and mechanisms of drug import could inform development of novel methods for antimalarial delivery.

Our findings add to a growing literature of host proteins that are internalized by *P. falciparum* (22, 28, 37, 38). The mechanism by which this occurs, and whether this represents a novel functional pathway or mis-trafficking, remains unknown. Importantly, this phenomenon is not observed for hemoglobin, the most abundant erythrocyte protein, suggesting that internalization may depend on specific protein features (37, 39). Moreover, not all internalized host proteins show the same localization: some are found in endocytic vesicles, while others, like APEH, appear diffusely within the parasite cytoplasm. These findings raise key questions about how host proteins might enter the parasite but avoid proteolytic digestion in the food vacuole. Elucidation of these mechanisms could provide novel insights into parasite biology and host-parasite interactions, and potentially unveil new therapeutic targets or methods of parasite-specific drug delivery.

## Materials and Methods

### P. falciparum culture

*P. falciparum* strain 3D7 (wild-type WT) was obtained through the MR4 as part of the BEI Resources Repository, NIAID, NIH (www.mr4.org). Parasites were cultured in a 2% hematocrit suspension of human erythrocytes in culture media (RPMI 1640 medium [Sigma] supplemented with 27 mM sodium bicarbonate, 11 mM glucose, 5 mM HEPES, 1 mM sodium pyruvate, 0.37 mM hypoxanthine, 0.01 mM thymidine, 10 µg/mL gentamicin, and 0.5% Albumax [Gibco]) at 37°C, 5% O_2_/5% CO_2_/90% N_2_ atmosphere as previous described (1).

### POM-ERJ, ERJ, POM-HEX, Fosmidomycin, and AA74-1

POM-ERJ (RCB-185) and ERJ were prepared as previously described (1). Fosmidomycin (Invitrogen) and AA74-1 (Sigma) were obtained from commercial vendors.

### Synthesis of fluorescent promoieties

Fluorescent promoieties were synthesized as previously described (23, 24, 31, 40–43). Synthesis and validation of additional promoieties are outlined in the **Supplemental Methods** and **Supplemental Appendix 1**.

### Native in-gel fluorescence activity

*P. falciparum* parasites were separated from erythrocytes using saponin. Saponin separated parasites and uninfected erythrocytes were lysed by sonication (Supplemental Methods). Lysate was separated by native polyacrylamide gel electrophoresis. Samples were mixed with 4x loading buffer containing 75 mM BisTris, 8 mM 6-AHA, 10% glycerol and 0.04% ponceau S and separated on NativePAGE™ 4 to 16% Bis-Tris gels (Invitrogen) with cathode buffer containing 50 mM tricine and 15 mM BisTris, and anode buffer containing 50 mM BisTris (pH 7.0). After protein separation, gels were incubated in Dulbecco’s phosphate buffered saline (dPBS) with 1.25 mM MgCl_2_ and 10 µM fluorescent substrate. Gels were imaged prior to incubation and at 30 minutes using a ChemiDoc MP imaging system (Bio-Rad).

### Peptide mass-spectroscopy

Fluorescent bands from native in-gel fluorescence assays were excised and cut into 1 mm^3^ cubes and subjected to in gel trypsin digestion. Resulting peptides were de-salted, dried by vacuum centrifugation and reconstituted in 0.1% TFA containing iRT peptides (Biognosys Schlieren, Switzerland). Peptides were analyzed on a QExactive HF mass spectrometer (ThermoFisher Scientific San Jose, CA) coupled with an Ultimate 3000 nano UPLC system and an EasySpray source using data dependent acquisition (DDA). The raw MS files were processed with MS Fragger and results loaded into Scaffold 5 for visualization.

### APEH Immunoblotting

Proteins from native in-gel fluorescence assays were transferred to PVDF membranes, blocked with phosphate buffered saline with 0.05% tween (PBST) and 5% bovine serum albumin (BSA), and probed with primary antibody (rabbit anti-APEH [Prestige Antibodies, HPA029700, Sigma]) diluted 1:5,000 in PBST with 5% BSA. Membranes were washed in PBST, and probed with secondary antibody (horseradish peroxidase tagged goat anti-rabbit [65-6120, Invitrogen]) diluted 1:20,000 in PBST with 5% BSA. Blots were imaged using SuperSignal™ West Pico PLUS Chemiluminescent Substrate (Thermo) on the ChemiDoc MP imaging system (Bio-Rad).

### APEH Immunofluorescence

*P. falciparum* infected erythrocytes were fixed with paraformaldehyde and glutaraldehyde, blocked with PBST + 3% BSA for 1 hour, and probed with primary antibody (rabbit anti-APEH [Prestige Antibodies, HPA029700, Sigma] or [PA5-97467, Invitrogen]) at dilutions from 1:100 to 1:900 in PBST + 3% BSA for 1 hour at room temperature. Cells were washed with PBST, probed with secondary antibody (Goat anti-Rabbit IgG (H+L) Cross-Adsorbed Secondary Antibody, Alexa Fluor™ 488, A-11008, ThermoFisher) for 1 hour at room temperature, and stained with 2 µM Hoechst for 10 minutes. Cells were imaged using a Nikon Eclipse Ti2 microscope in Poly-D-Lysine (Thermo) coated µ-Plate 96 well square plates (Ibidi).

### Measurement of POM-ERJ activation *in vitro* via DXR activity assay

6-HIS tagged recombinant *Escherichia coli* DXR was expressed in *E. coli* cells and purified using Nickel-NTA resin (Qiagen) followed by size exclusion chromatography as previously described (9). 10 µM POM-ERJ was diluted in assay buffer containing 25 mM Tris (pH 7.5), 100 mM NaCl, 7.5 mM MgCl_2_, 0.1 mg/ml BSA and incubated with *P. falciparum* lysate (80 ng/ul), erythrocyte lysate (80 ng/ul), or rAPEH (2 ng/ul) in the presence or absence of 100 nM AA74-1 for 12 hours at 37 °C. Recombinant DXR and NADPH were then added to each reaction, followed by 1-deoxy-D-xylulose 5-phosphate (DOXP), and DXR activity measured as previously described (44). For the Fsm condition, 10 µM Fsm was incubated for 12 hours in assay buffer prior to mixing with DXR and NADPH.

### Activation of POM-ERJ measured by mass spectrometry

Reactions containing dPBS with 2 mM MgCl_2_, 10 µM POM-ERJ, and either 15 ng/µl rAPEH, 15 ng/µl rAPEH and 400 nM AA74-1, or buffer were incubated at 37 °C and sampled at 0, 1, 2, 4, and 6 hours and analyzed as previously described (11), with modifications (Supplemental Materials and Methods).

### *P. falciparum* EC_50_ for POM-ERJ and Fsm with or without AA74-1 pretreatment

Erythrocytes in culture media were incubated with 1 µM AA74-1 or DMSO at 37 °C for 24 hours and then washed three times in culture media. Cultures of *P. falciparum* strain 3D7 were grown in either AA74-1 treated (APEH inhibited) or DMSO treated (APEH active) erythrocytes for at least 1 week prior to growth inhibition analyses to remove the majority of untreated erythrocytes. Cultures were then diluted to 1% parasitemia in either AA74-1 pretreated (APEH inhibited) or DMSO pretreated (APEH active) erythrocytes and treated with POM-ERJ (1.95 nM to 1 µM) or Fsm (48 nM to 25 µM) for three days. Parasite growth was quantified and EC_50_ values calculated as described above.

### Measurement of POM-HEX activation via erythrocyte glycolysis

Erythrocytes were pretreated with 400 nM AA74-1 or DMSO at 37 °C for 24 hours, washed 3x with dPBS and resuspended at 2% hematocrit in Seahorse XF DMEM assay medium (pH 7.4) (Agilent) containing 10 mM glucose (Agilent). Erythrocytes were transferred to a Poly-D-Lysine (Thermo) coated Seahorse XF 96 well microplate (200 µl per well) and centrifuged at 1000 x g for 10 minutes. Extracellular acidification rates were monitored using the Agilent Seahorse XFe96. After the third cycle, 10 µM POM-HEX in DMEM, or DMEM alone, was added to the wells.

### Activation of fluorescent promoieties by rAPEH and parasite lysate

Fluorescent promoieties (22 µM) were diluted in dPBS with 2 mM MgCl_2_ and mixed with rAPEH (1.45 ng/ul), parasite lysate (160-189 ng/ul), or parasite lysate with 250 nM AA74-1 in low volume 384-well black flat bottom microplates (Corning). Change in fluorescence (λ_ex_ = 485 nm, λ_em_ = 520 nm) was monitored on a FLUOstar Omega microplate reader (BMG Labtech) at 37°C. Fluorescence measurements were converted to molar concentrations using a fluorescein standard curve. Initial rates of reactions were calculated, corrected for the rate of background hydrolysis in buffer alone, and normalized by protein concentration (measured by Qubit protein assay [ThermoFisher]). Calculation of reaction rates and subsequent analyses were performed in R (45).

## Acknowledgements

Peptide mass spectrometry was performed by the Children’s Hospital of Philadelphia Research Institute Proteomics Core Facility (RRID:SCR_023099). Mass spectrometry of POM-ERJ was performed by the Small Molecule and Metabolite Core. We thank Dr. Weimin Liu and Dr. Beatrice Hahn for providing 293F cell cultures and media for APEH expression. Need to add grant numbers here.

---

**Figure S1:**
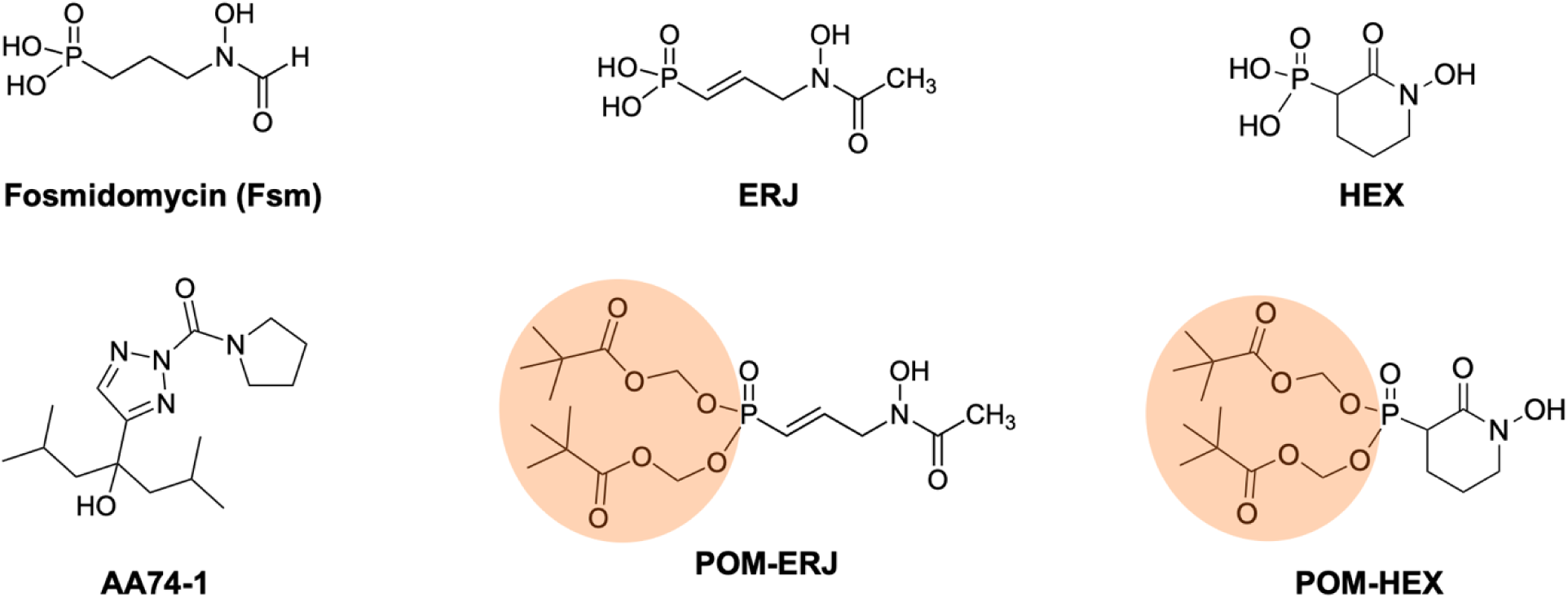
Chemical structures of inhibitors and prodrugs used in this study. Shown are the DXR inhibitors (Fsm and ERJ), the pivaloyloxymethyl prodrug of ERJ (POM-ERJ), the enolase inhibitor (HEX), the pivaloyloxymethyl prodrug of HEX (POM-HEX), and the APEH inhibitor (AA74-1). Pivaloyloxymethyl prodrug moieties are highlighted in orange.

**Figure S2:**
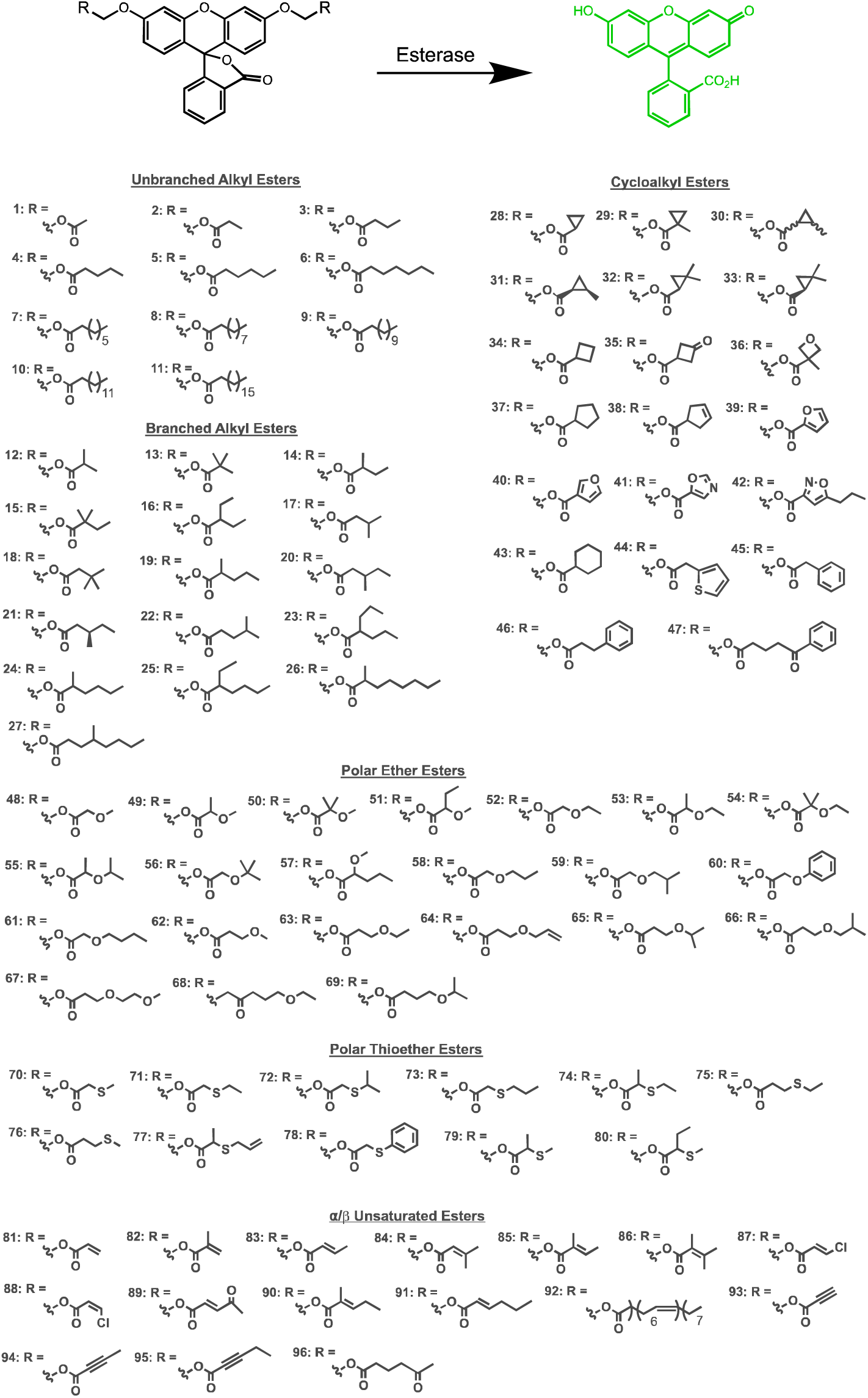
Full substrate library used in this study. Substrates are divided by type: branched alkyl, unbranched alkyl, cycloalkyl, polar ether, polar thioether, and α/β unsaturated esters.

**Figure S3.**
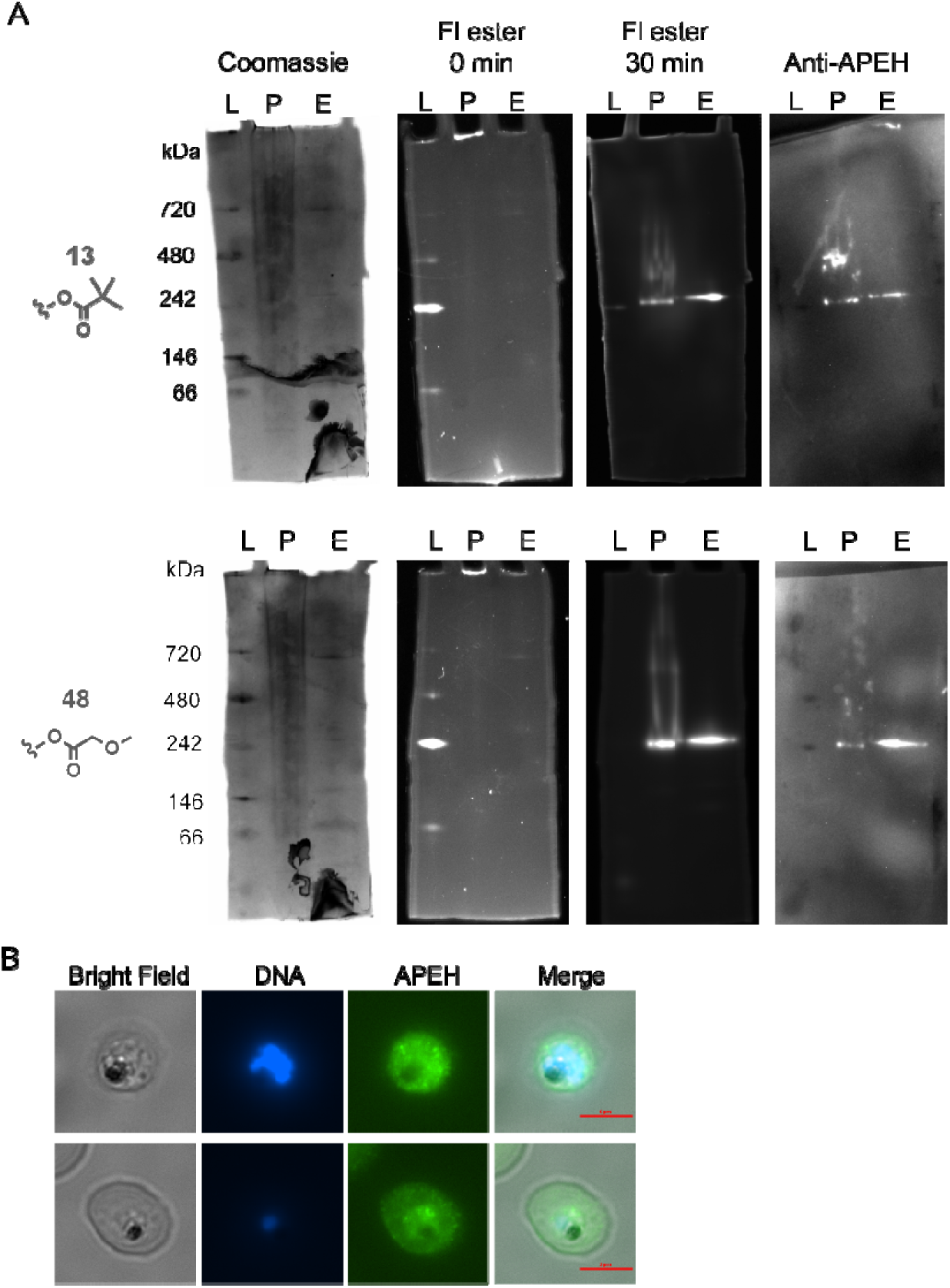
Parasite localized APEH activates fluorescent promoieties and localizes to the parasite cytoplasm. **A**) Full images of in-gel fluorescent activity assays and immunoblotting for compounds 13 and 48. Presented from left to right: Coomassie stained gel, background fluorescence before incubation with fluorescent ester (0 min), fluorescence after 30-minute incubation with substrate, and immunoblot with APEH (Anti-APEH). Ladder proteins that migrate at 480, 242, and 66 kDa exhibit slight fluorescence. *L: ladder, P: P. falciparum lysate, E: erythrocyte lysate*. **B**) Indirect immunofluorescence images using a second anti-APEH antibody (Prestige Antibodies, Sigma).

**Figure S4.**
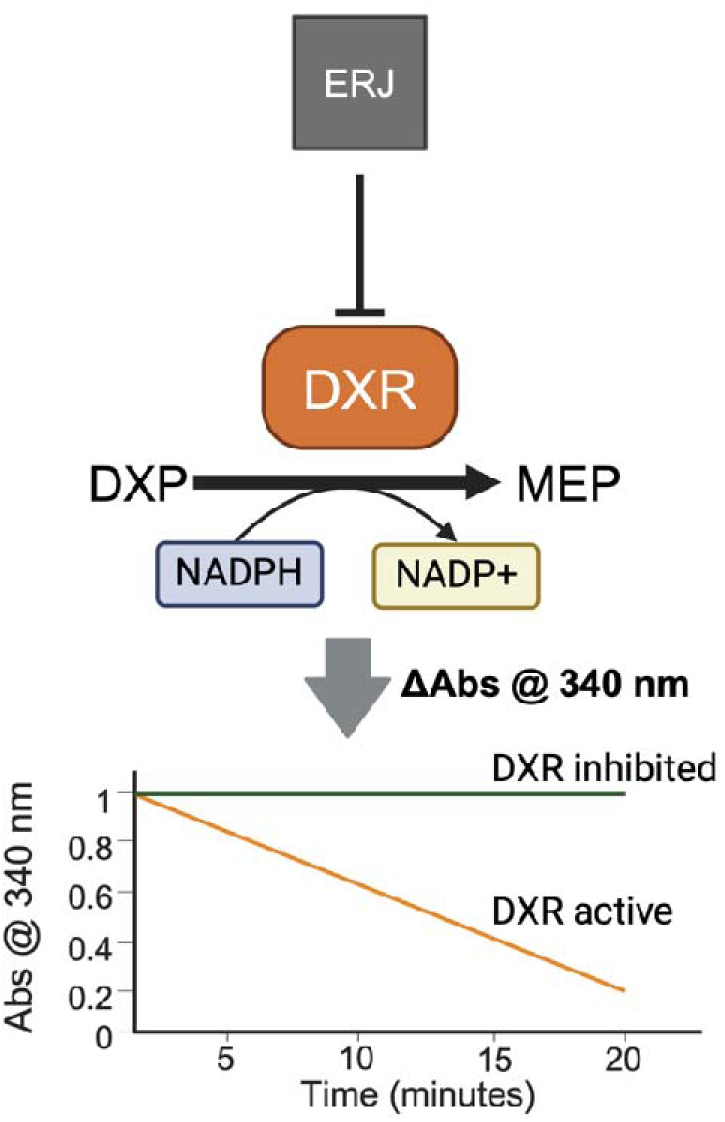
Schematic of DXR activity assay. DXR converts 1-deoxy-D-xylulose 5-phosphate (DXP) to 2-C-methyl-D-erythritol 4-phosphate (MEP), converting NADPH to NADP^+^ in the process. The concentration of NADPH is measured by the absorbance at 340 nm (Abs @ 340 nm), with a decline in absorbance indicating conversion of NADPH to NADP^+^ by active DXR.

**Figure S5.**
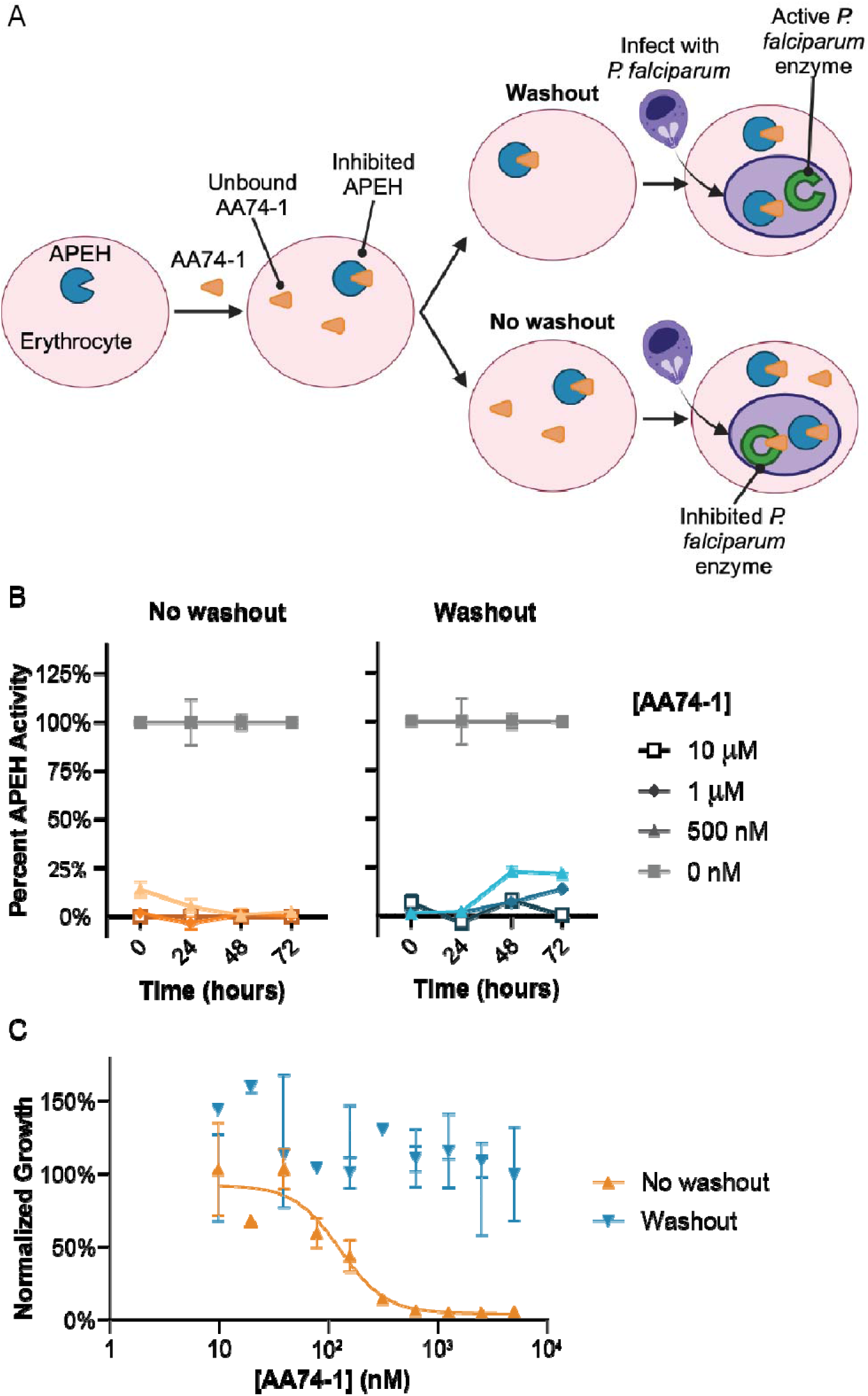
Pretreatment of erythrocytes with AA74-1 followed by washout leads to sustained APEH inhibition but restores parasite killing. **A**) Schematic of washout growth inhibition assay for AA74-1. Treatment of erythrocytes with excess AA74-1 leads to irreversible inhibition of APEH and residual unbound AA74-1. Without washout, when erythrocytes are infected with *P. falciparum*, this unbound AA74-1 can inhibit additional *P. falciparum* enzymes. Washout removes unbound AA74-1, leading to decreased inhibition of additional *P. falciparum* enzymes but sustained inhibition of APEH. **B**) Residual APEH activity, quantified by rates of acetylalanine 4-nitroanilide hydrolysis, in uninfected erythrocytes after AA74-1 treatment, with or without washout. **C**) Dose response curves for *P. falciparum* 3D7 growth in erythrocytes pretreated with AA74-1 with or without washout. Erythrocytes were incubated with AA74-1 for 24 hours prior to infection. Erythrocytes were incubated with AA74-1 for 24 hours prior to the start of the APEH activity assay or 72-hour growth inhibition assay. For washed replicates, erythrocytes were washed three times in media. Data represent mean ± SEM of two technical replicates.

**Figure S6.**
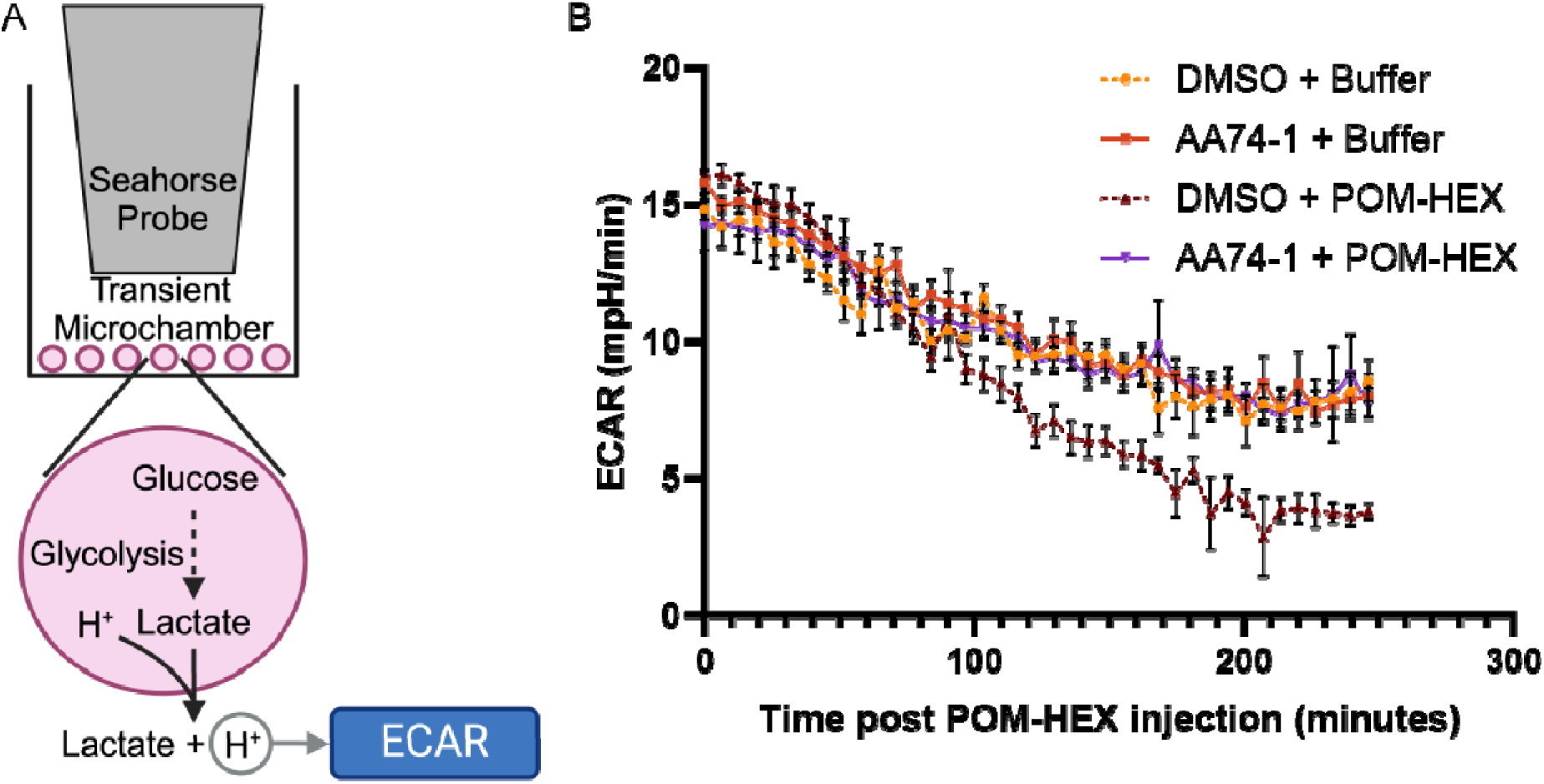
APEH activates POM-HEX in uninfected erythrocytes. A) Schematic of the Agilent Seahorse XF. Lowering the assay probe creates a transient microchamber. Glycolysis results in the production of lactate, which is transported out of the cell along with a proton. The rate of extracellular acidification (ECAR) is thus a proxy for the rate of glycolysis. **B**) ECAR of erythrocytes treated with POM-HEX or control (buffer) after pre-incubation with AA74-1 or DMSO for 24 hours. Addition of POM-HEX decreases glycolysis relative to control erythrocytes, but this decrease is not seen in erythrocytes pretreated with AA74-1. Data (mean ± SEM) are representative of 4 technical replicates. *mpH/min: mili pH per minute*

**Figure S7.**
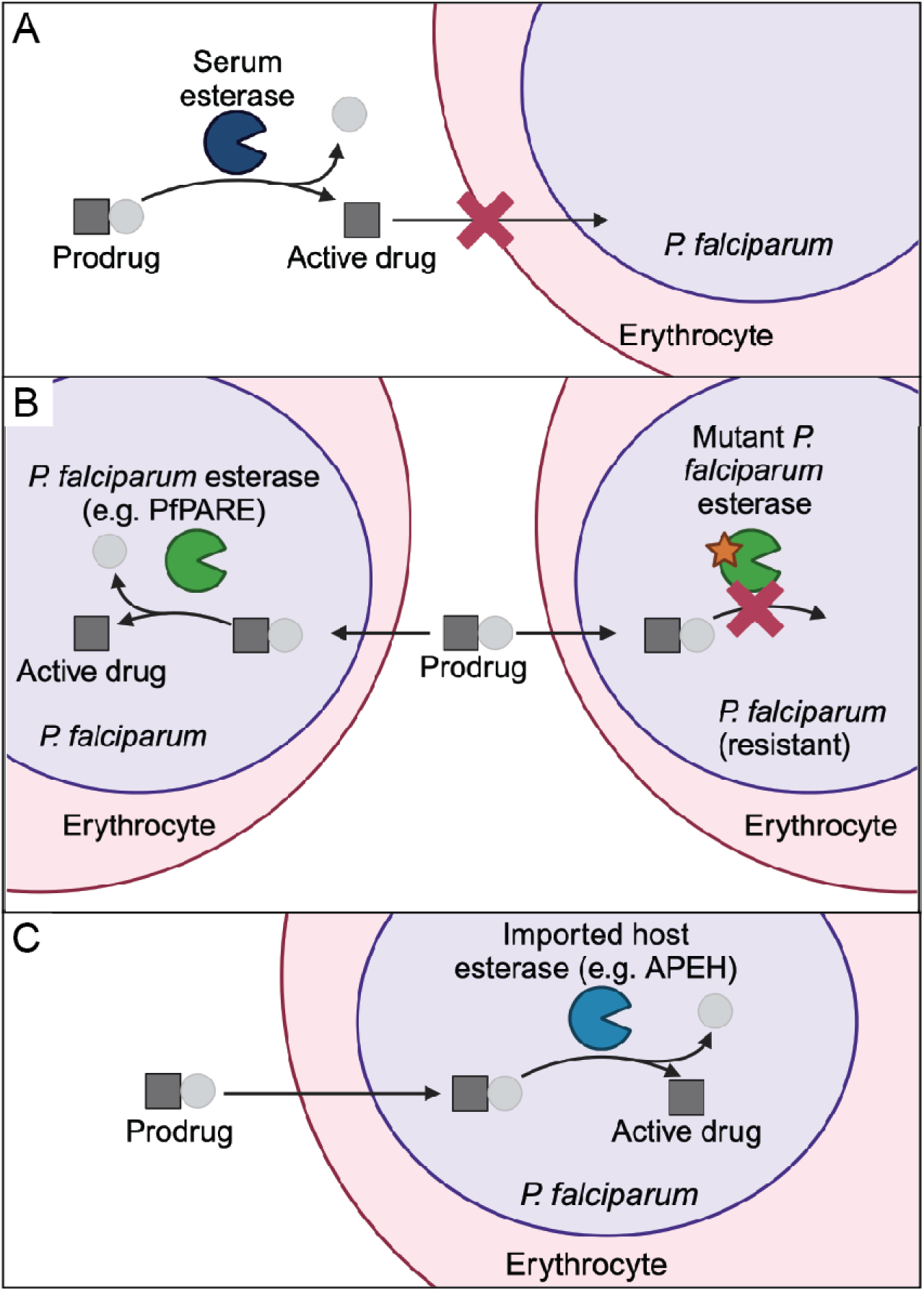
Prodrug activation by serum, *P. falciparum*, or imported host esterases. **A**) Prodrugs activated by serum esterases lose beneficial prodrug properties (*i.e.*, improved cellular permeability) prior to reaching their target, decreasing accumulation at the target site. **B**) Prodrugs activated exclusively by *P. falciparum* esterases are not activated in the serum, maintaining prodrug properties until activated within the parasite. However, mutations in the prodrug activation enzyme may lead to resistance. PfPARE: *P. falciparum* Prodrug Activation and Resistance Esterase. **C**) Prodrugs activated by imported host esterases maintain prodrug properties until reaching their target. As these prodrugs are activated by host enzymes, resistance is less likely to develop through mutations in activating enzymes.

