## Supplemental Methods for "Prodrug activation in malaria parasites mediated by an imported erythrocyte esterase, acylpeptide hydrolase (APEH)"

Synthesis of fluorescent promoieties

The synthesis of compounds (**3,4,5)**^1^, (**1,2,13,29,34,82**)^2^, (**37,39,43,45,48,62,81**)^3^, (**9,11**)^4^, (**16,17,18,19,20,22,25,35,50,58,90**)^5^, (**14,15,46,49,51,52,53,60,63,70,71,72,73,74,75,76,77,78,79,80**)^6^, and (**12,28,32,41,67,83,94**)^7^, has been previously described . Unless otherwise noted, all chemicals were purchased from Sigma Aldrich and used without further purification. Carboxylic acid reagent used for the synthesis of compounds 23, 27, 61, 84, 87, 88, 89, 92, 93, and 96 were from AlfaAesar. Carboxylic acid reagent used for the synthesis of compounds 24, 65, 66, 68, 69, 86, 91, and 95 were from Enamine. Carboxylic acid reagent used for the synthesis of compounds 33, 36, 49, and 44 were from Combiblocks. Carboxylic acid reagent used for the synthesis of compounds 54, 55, and 57 were from Matrix. Carboxylic acid reagent used for the synthesis of compound 64 was from Ambeed. Carboxylic acid reagent used for the synthesis of compound 85 was from Oakwood. All reactions were monitored using Macherey-Nagel analytical thin layer chromatography (TLC) plates (POLYGRAM® SIL G/UV_254_, polyester back). Nuclear magnetic resonance (NMR) spectra were obtained using a Bruker Biospin Avance III HD 400 operating at 400.19 MHz for ^1^H and 100.64 MHz for ^13^C. High resolution mass spectrometry (HRMS) was performed with electrospray ionization (ESI) by the Mass Spectrometry facility at the Department of Chemistry, Indiana University using an Agilent 1200 HPLC-6130 MSD mass spectrometer **(Supplemental Appendix 1)**.

The following procedure is representative for the synthesis of all new compounds, using Fluorescein bis(3-methylbutanyloxymethyl ether)^5^ as a starting material.

*Synthesis of Fluorescein bis(3-methylbutanyloxymethyl ether)* (**6**). Fluorescein bis(chloromethyl ether) (30.0 mg, 69.9 μmol, 1.0 Eq), heptanoic acid (36.3 mg, 279.6 μmol, 4.0 Eq.), and Cs_2_CO_3_ (91.1 mg, 279.6 μmol, 4.0 Eq.) were dissolved in dry CH_3_CN (1 mL). Molecular sieves (100 mg) were added, and the reaction was covered in foil and allowed to stir for 24 h at ambient temperature. The reaction mixture was adsorbed onto celite and purified via column chromatography using a hexane to ethyl acetate gradient in solvent containing constant 20% dichloromethane.

Characterization data for all new compounds

*Fluorescein bis*(*heptanyloxymethyl ether)*

Data for **6**: (69%, white solid).  ^1^H-NMR (CDCl_3_, 400 MHz): δ = 8.03 (d, 1H), 7.7-7.6 (m, 2H), 7.15 (d, 1H), 6.96 (m, 2H), 6.73 (m, 4H), 5.79 (s, 4H), 2.37 (t, 4H), 1.67-1.60 (m, 4H), 1.32-1.23 (m, 12H), 0.85 (t, 6H) ppm.  ^13^C-NMR (CDCl_3_, 100 MHz): δ = 172.6, 169.2, 158.4, 153.0, 152.2, 135.1, 129.9, 129.4, 126.7, 125.2, 123.9, 113.2, 112.7, 103.5, 84.8, 82.4, 34.2, 31.4, 28.7, 24.6, 22.4, 14.0 ppm. HRMS (APCI): calc’d for MH^+^ = C_36_H_40_O_9_H: 617.2745, found: 617.2744.

*Fluorescein bis(octanyloxymethyl ether)*

Data for **7**: (30%, white solid). ^1^H-NMR (CDCl_3_, 400 MHz): δ = 8.03 (d, 1H), 7.7-7.6 (m, 2H), 7.15 (d, 1H), 6.96  (s, 2H), 6.73 (s, 4H), 5.79 (s, 4H), 2.37 (t, 4H), 1.7-1.6 (m, 4H), 1.3-1.2 (m, 16H), 0.84 (t, 6H)  ppm.  ^13^C-NMR (CDCl_3_, 100 MHz): δ = 172.6, 169.2, 158.4, 152.9, 152.2, 135.1, 129.9, 129.4, 126.6, 125.2, 123.8, 113.2, 112.7, 103.5, 84.8, 82.4, 36.6, 34.1, 31.6, 28.9, 24.7, 22.6, 14.0 ppm. HRMS (ESI): calc’d for MH^+^ C_42_H_52_O_9_H: 645.3063; found: 645.3037.

*Fluorescein bis*(*decanyloxymethyl ether)*

Data for **8**: (21%, white solid).  ^1^H-NMR (CDCl_3_, 400 MHz): δ = 8.05 (d, 1H), 7.7-7.6 (m, 2H), 7.17 (d, 1H), 6.98 (s, 2H), 6.76 (s, 4H), 5.81 (s, 4H), 2.40 (t, 4H), 1.7-1.6 (m, 4H), 1.3-1.2 (m, 24H), 0.88 (t, 6H) ppm.  ^13^C-NMR (CDCl_3_, 100 MHz): δ = 172.6, 169.2, 158.4, 153.0, 152.2, 135.1, 129.9, 129.4, 126.6, 125.2, 123.8, 113.2, 112.7, 103.5, 84.8, 82.4, 34.2, 31.8, 29.4, 29.2, 29.2, 29.0, 24.7, 22.6, 14.1 ppm. HRMS (ESI): calc’d for MH^+^ C_42_H_52_O_9_H: 701.3690; found: 701.3688.

*Fluorescein bis*(*tetradecanyloxymethyl ether)*

Data for **10**: (61%, white solid).  ^1^H-NMR (CDCl_3_, 400 MHz): δ = 8.05 (d, 1H), 7.7-7.6 (m, 2H), 7.17 (d, 1H), 6.98 (s, 2H), 6.76 (d, *J* = 1.4 Hz, 4H), 5.81 (s, 4H), 2.40 (t, 4H), 1.7-1.6 (m, 4H), 1.3-1.2 (m, 40H), 0.90 (t, 6H) ppm.  ^13^C-NMR (CDCl_3_, 100 MHz): δ = 172.6, 169.2, 158.4, 153.0, 152.2, 135.1, 129.9, 129.4, 126.6, 125.2, 123.8, 113.2, 112.7, 103.5, 84.8, 82.4, 34.1, 31.9, 29.7, 29.6, 29.6, 29.4, 29.4, 29.3, 29.2, 29.0, 24.7, 22.7, 14.1 ppm. HRMS (ESI): calc’d for MH^+^ C_50_H_68_O_9_H: 813.4941; found: 813.4944.

*Fluorescein bis((3R)-3-methylpentanyloxymethyl ether)*

Data for **21**: (75%, white solid).  ^1^H-NMR (CDCl_3_, 400 MHz): δ = 8.03 (d, 1H), 7.7-7.6 (m, 2H), 7.15 (d, 1H), 6.96 (m, 2H), 6.73 (m, 4H), 5.79 (s, 4H), 2.40-2.35 (dd, 2H), 2.21-2.16 (dd, 2H), 1.94-1.84 (m, 2H), 1.40-1.18 (m, 4H), 0.92 (d, 6H), 0.86 (t, 6H) ppm.  ^13^C-NMR (CDCl_3_, 100 MHz): δ = 172.1, 169.2, 158.4, 153.0, 152.2, 135.1, 129.9, 129.4, 126.6, 125.2, 123.9, 113.2, 112.7, 103.5, 84.7, 82.5, 41.2, 31.8, 29.2, 19.2, 11.2 ppm. HRMS: calc’d for MH^+^ = C_34_H_36_O_9_H: 589.2432, found: 589.2436.

*Fluorescein bis*((*2-(1-propyl*))*pentanyloxymethyl ether)* (presumably a racemic mixture)

Data for **23**: (29%, white solid).  ^1^H-NMR (CDCl_3_, 400 MHz): δ = 8.03 (d, 1H), 7.7-7.6 (m, 2H), 7.13 (d, 1H), 6.95 (m, 2H), 6.73 (m, 4H), 5.79 (s, 4H), 2.44-2.40 (m, 2H), 1.62-1.55 (m, 4H), 1.47-1.42 (m, 8H), 1.41-1.33 (sex, 8H), 0.83 (t, 12H) ppm.  ^13^C-NMR (CDCl_3_, 100 MHz): δ = 175.2, 169.3, 158.4, 153.1, 152.2, 135.1, 129.9, 129.3, 129.2, 126.6, 125.2, 123.8, 113.2, 112.7, 103.6, 84.7, 82.4, 45.2, 44.6, 34.3, 20.5, 13.9 ppm. HRMS (ESI): calc’d for MNa^+^ C_38_H_44_O_9_Na: 667.2878; found: 667.2879.

*Fluorescein bis(2-methylhexanyloxymethyl ether)* (presumably a racemic mixture)

Data for **24**: (53%, white solid).  ^1^H-NMR (CDCl_3_, 400 MHz): δ = 8.03 (d, 1H), 7.7-7.6 (m, 2H), 7.14 (d, 1H), 6.96 (m, 2H), 6.74 (m, 4H), 5.82-5.77 (m, 4H), 2.52-2.47 (m, 2H), 1.68-1.63 (m, 2H), 1.46-1.42 (m, 2H) 1.25-1.23 (m, 8H), 1.16 (d, 6H), 0.81 (t, 6H) ppm.  ^13^C-NMR (CDCl_3_, 100 MHz): δ = 175.6, 169.2, 158.4, 152.9, 152.2, 135.0, 129.8, 129.3, 126.6, 125.1, 123.8, 113.1, 112.7, 103.4, 84.7, 82.4, 39.4, 33.2, 29.2, 22.5, 16.8, 13.8 ppm. HRMS: calc’d for MH^+^ = C_36_H_40_O_9_H: 617.2745, found: 617.2747.

*Fluorescein bis(2-methylheptanyloxymethyl ether)* (non-racemic mixture)

Data for **26**: (34%, white solid).  ^1^H-NMR (CDCl_3_, 400 MHz): δ = 8.05 (d, 1H), 7.7-7.6 (m, 2H), 7.17 (d, 1H), 6.98 (t, 2H), 6.75 (d, 4H), 5.81 (s, 3H), 2.57-2.48 (m, 2H), 1.70-1.62 (m, 2H), 1.49-1.41 (m, 2H), 1.35-1.15 (m, 20H (includes a doublet at 1.18 ppm)), 0.84 (t, 6H) ppm. ^13^C-NMR (CDCl_3_, 100 MHz): δ = 175.6, 169.2, 158.5, 153.0, 152.2, 135.1, 129.9, 129.3, 126.7, 125.2, 123.8, 113.2, 112.7, 103.5, 84.8, 82.5, 39.5, 33.5, 31.6, 29.1, 27.0, 22.5, 16.8, 14.0 ppm (the C-NMR has both the peaks listed here for the major stereoisomer (ranging from 39.5 to 14.0 ppm), and slightly different peaks for the minor stereoisomer, indicating a non-racemic mixture – see spectra provided in Supplemental Information to see the detailed peaks associated with the acyl chain). HRMS: calc’d for MH^+^ = C_40_H_48_O_9_H: 673.3371, found: 673.3371.

*Fluorescein bis(4-methyloctanyloxymethyl ether)* (presumably a racemic mixture)

Data for **27**: (48%, white solid).  ^1^H-NMR (CDCl_3_, 400 MHz): δ = 8.03 (d, 1H), 7.7-7.6 (m, 2H), 7.15 (d, 1H), 6.96 (m, 2H), 6.73 (m, 4H), 5.79 (s, 4H), 2.41-2.35 (m, 4H), 1.71-1.65 (m, 2H), 1.48-1.38 (m, 4H), 1.28-1.17 (m, 13H), 0.88-0.84 (overlapping t and d, 13H) ppm. ^13^C-NMR (CDCl_3_, 100 MHz): δ = 172.9, 169.2, 158.4, 153.0, 152.2, 135.1, 129.9, 129.4, 126.6, 125.2, 123.9, 113.2, 112.7, 103.5, 84.8, 82.5, 36.3, 32.3, 31.9, 31.6, 29.1, 22.9, 19.2, 14.1 ppm. HRMS (APCI): calc’d for MH^+^ = C_40_H_48_O_9_H: 673.3371, found: 673.3374; calc’d for MNa^+^ C_40_H_48_O_9_Na: 695.3191, found: 695.3192.

*Fluorescein bis*(*2-methylcyclopropane-1-carboxymethyl ether)* (mixture of several stereoisomers)

Data for **30**: (36%, white solid).  ^1^H-NMR (CDCl_3_, 400 MHz): δ = 8.03 (d, 1H), 7.7-7.6 (m, 2H), 7.16 (d, 1H), 6.98 (m, 2H), 6.74 (s, 4H), 5.80-5.75 (m, 4H), 1.75-1.70 (m, 1H), 1.50-1.45 (m, 2H), 1.41-1.37 (m, 2H), 1.27-1.22 (m, 2H), 1.18 & 1.12 (d’s, 6H), 0.98-0.94 (m, 1H), 0.85-0.80 (m, 2H), 0.79-0.74 (m, 1H)  ppm.  ^13^C-NMR (CDCl_3_, 100 MHz): δ = 173.3, 171.8, 169.2, 158.52, 158.48, 153.0, 152.3, 135.1, 129.9, 129.4, 126.7, 125.2, 123.9, 113.2, 112.7, 103.6, 103.5, 84.9, 84.7, 82.5, 21.1, 18.5, 18.3, 17.8, 17.6, 17.0, 15.3, 12.1 ppm. HRMS (ESI): calc’d for MH^+^ = C_32_H_29_O_9_: 557.1806, found: 557.1805; calc’d for MNa^+^ C_32_H_28_O_9_Na: 579.1626, found: 579.1625.

*Fluorescein bis*(*cis-2-methylcyclopropane-1-carboxymethyl ether)* (mixture of stereoisomers, but all *cis*)

Data for **31**: (48%, white solid).  ^1^H-NMR (CDCl_3_, 400 MHz): δ = 8.03 (d, 1H), 7.7-7.6 (m, 2H), 7.16 (d, 1H), 6.98 (m, 2H), 6.74 (m, 4H), 5.80 (m, 4H), 1.75-1.69 (m, 2H), 1.40-1.34 (m, 2H), 1.18 (d, 6H), 1.13-1.07 (m, 2H), 0.98-0.94 (m, 2H) ppm.  ^13^C-NMR (CDCl_3_, 100 MHz): δ = 171.8, 169.3, 158.5, 153.0, 152.2, 135.1, 129.9, 129.3, 126.7, 125.1, 123.9, 113.1, 112.8, 103.6, 103.5, 84.7, 82.5, 18.4, 17.1, 15.3, 12.1 ppm. HRMS: calc’d for MH^+^ = C_32_H_28_O_9_H: 557.1806, found: 557.1807.

*Fluorescein bis((1S)-2,2-dimethylcyclopropane-1-carboxymethyl ether)*

Data for **33**: (63%, white solid).  ^1^H-NMR (CDCl_3_, 400 MHz): δ = 8.03 (d, 1H), 7.7-7.6 (m, 2H), 7.17 (d, 1H), 6.98 (m, 2H), 6.74 (m, 4H), 5.83-5.75 (dd, 4H), 1.57-1.54 (m, 2H), 1.22 (s, 6H), 1.15-1.17 (m, 8H), 0.96-0.93 (m, 2H) ppm.  ^13^C-NMR (CDCl_3_, 100 MHz): δ = 171.6, 169.1,158.4, 152.8, 152.1, 135.0, 129.7, 129.2, 126.5, 125.0, 123.7, 113.0, 112.9, 112.7, 112.6, 103.4, 103.3, 84.6, 82.4, 26.7, 26.4, 24.3, 22.8, 18.6 ppm. HRMS: calc’d for MH^+^ = C_34_H_32_O_9_H: 585.2119, found: 585.2123.

*Fluorescein bis(2-(2-oxetanyl)propanyloxymethyl ether)*

Data for **36**: (66%, white solid).  ^1^H-NMR (CDCl_3_, 400 MHz): δ = 8.06 (d, 1H), 7.73-7.65 (m, 2H), 7.20 (d, 1H), 6.99 (s, 2H), 6.77 (s, 4H), 5.89 (s, 4H), 4.96 (d, 4H), 4.44 (dd, 4H), 1.65 (s, 6H) ppm.  ^13^C-NMR (CDCl_3_, 100 MHz): δ = 173.2, 169.1, 158.2, 152.8, 152.2, 135.2, 129.9, 129.5, 126.6, 125.2, 123.9, 113.6, 112.6, 103.6, 100.0, 85.5, 79.2, 44.6, 21.6 ppm. HRMS (ESI): calc’d for MH^+^ C_32_H_28_O_11_H: 589.1710; found: 589.1730.

*Fluorescein bis*(*cyclopentene-4-carboxymethyl ether)*

Data for **38**: (39%, white solid).  ^1^H-NMR (CDCl_3_, 400 MHz): δ = 8.05 (d, 1H), 7.73-7.64 (m, 2H), 7.19 (d, 1H), 6.99 (t, 2H), 6.76 (d, 4H), 5.83, (s, 4H), 5.68 (s, 4H), 3.21 (m, 2H), 2.69 (d, *J* = 8.1 Hz, 8H) ppm.  ^13^C-NMR (CDCl_3_, 100 MHz): δ = 174.9, 169.2, 158.5, 152.9, 152.2, 135.1, 129.9, 129.4, 128.8, 126.7, 125.2, 123.9, 113.3, 112.7, 103.6, 85.1, 82.5, 41.4, 36.1 ppm. HRMS (ESI): calc’d for MH^+^ C_34_H_28_O_9_H: 581.1812; found: 581.1795.

*Fluorescein bis (furan-3-carboxymethyl ether)*

Data for **40**: (14%, white solid).  ^1^H-NMR (CDCl_3_, 400 MHz): δ = 8.08 (s, 2H), 8.02 (d, 1H), 7.67-7.63 (m, 2H), 7.45 (s, 2H), 7.15 (d, 1H), 7.01 (d, 2H), 6.78-6.75 (m, 6H), 5.96 (s, 4H) ppm.  ^13^C-NMR (CDCl_3_, 100 MHz): δ = 169.2, 161.6, 158.4, 152.9, 152.2, 148.6, 144.1, 135.1, 129.9, 129.4, 126.6, 125.2, 123.9, 118.4, 113.4, 112.7, 109.8, 103.6, 85.1, 82.4 ppm. HRMS (ESI): calc’d for MH^+^ C_32_H_20_O_11_H: 581.1084; found: 581.1085.

*Fluorescein bis(((5-n-propyl)isoxazole-3-carboxy)methyl ether)*

Data for **42**: (42%, white solid).  ^1^H-NMR (CDCl_3_, 400 MHz): δ = 8.01 (d, 1H), 7.68-7.59 (m, 2H), 7.12 (d, 1H), 7.03 (d, 2H), 6.81-6.72 (m, 4H), 6.44 (s, 2H), 6.02 (s, 4H), 2.77 (t, 4H), 1.79-1.69 (m, 4H), 0.98 (t, 6H) ppm.  ^13^C-NMR (CDCl_3_, 100 MHz): δ = 176.0, 169.2, 159.1, 158.1, 155.4, 152.9, 152.1, 135.1, 129.9, 129.5, 126.4, 125.1, 123.9, 113.6, 112.7, 103.8, 101.7, 86.0, 82.2, 28.6, 20.8, 13.5 ppm. HRMS (APCI): calc’d for MH^+^ = C_36_H_30_O_11_N_2_H: 667.1922, found: 667.1924; calc’d for MNa^+^ = C_36_H_30_O_11_N_2_Na: 689.1742, found: 689.1744.

*Fluorescein bis((2-thiophene)acetyloxymethyl ether)*

Data for **44**: (18%, white solid).  ^1^H-NMR (CDCl_3_, 400 MHz): δ = 8.07 (dm, 1H), 7.73-7.65 (m, 2H), 7.23-7.22 (m, 2H), 7.18 (dm, 1H), 6.97-6.96 (m, 6H), 6.76-6.71 (m, 4H), 5.85 (s, 4H), 3.94 (s, 4H) ppm. ^13^C-NMR (CDCl_3_, 100 MHz): δ = 169.3, 169.2, 158.2, 152.9, 152.2, 135.1, 133.8, 129.9, 129.4, 127.2, 126.9, 126.6, 125.4, 125.2, 123.9, 113.4, 112.7, 103.7, 85.5, 82.4, 35.3 ppm. HRMS: calc’d for MH^+^ = C_34_H_24_O_9_S_2_H: 641.0935, found: 641.0932; calc’d for MNa^+^ = C_34_H_24_O_9_S_2_Na: 663.0754, found: 663.0750.

*Fluorescein bis*((*5-oxo-5-phenyl)pentanyloxymethyl ether)*

Data for **47**: (36%, white solid).  ^1^H-NMR (CDCl_3_, 400 MHz): δ = 8.05 (d, 1H), 7.93 (d, 4H), 7.70-7.63 (m, 2H), 7.55 (t, 2H), 7.44 (t, 4H), 7.13 (d, 1H), 6.97 (s, 2H), 6.74 (s, 4H), 5.82, (s, 4H), 3.07 (t, 4H), 2.55 (t, 4H), 2.08-2.16 (m, 4H) ppm.  ^13^C-NMR (CDCl_3_, 100 MHz): δ = 199.1, 172.1, 169.2, 158.3, 152.9, 152.2, 136.7, 135.1, 133.2, 129.9, 129.4, 128.6, 128.0, 126.6, 125.1, 123.9, 113.3, 112.7, 103.5, 84.8, 82.4, 37.1, 33.2, 19.0 ppm. HRMS (ESI): calc’d for MH^+^ C_44_H_36_O_11_H: 741.2336; found: 741.2327.

*Fluorescein bis*((*2-ethoxy-2-methyl*)*propanyloxymethyl ether)*

Data for **54**: (57%, white solid).  ^1^H-NMR (CDCl_3_, 400 MHz): δ = 8.05 (d, 1H), 7.72-7.64 (m, 2H), 7.16 (d, 1H), 7.00 (m, 2H), 6.76 (s, 4H), 5.86 (s, 4H), 3.40 (q, 4H), 1.46 (s, 12 H), 1.14 (t, 6H) ppm.  ^13^C-NMR (CDCl_3_, 100 MHz): δ = 173.8, 169.2, 158.3, 152.9, 152.2, 135.1, 129.9, 129.4, 126.6, 125.2, 123.8, 113.4, 112.7, 103.6, 85.3, 82.3, 60.4, 24.6, 15.6 ppm. HRMS (ESI): calc’d for MH^+^ C_34_H_36_O_11_H: 621.2336; found: 621.2324.

*Fluorescein bis*((*2-isopropoxy*)*propanyloxymethyl ether)* (presumably racemic mixture)

Data for **55**: (45%, white solid).  ^1^H-NMR (CDCl_3_, 400 MHz): δ = 8.06 (d, 1H), 7.72-7.64 (m, 2H), 7.16 (d, 1H), 6.99 (s, 2H), 6.76 (s, 4H), 5.87 (dd, J = 6.6, 32.5 Hz, 4H), 4.13 (q, 2H), 3.63 (sep, 2H), 1.41 (d, 6H), 1.17 (t, 11H) ppm.  ^13^C-NMR (CDCl_3_, 100 MHz): δ = 172.9, 169.2, 158.3, 152.9, 152.2, 135.1, 129.9, 129.4, 126.6, 125.2, 123.8, 113.4, 112.8, 103.6, 85.1, 82.3, 72.1, 71.8, 22.8, 21.4, 19.1, 18.6 ppm. HRMS (ESI): calc’d for MH^+^ C_34_H_36_O_11_H: 621.2336; found: 621.2327.

*Fluorescein bis((t-butoxy)acetyloxymethyl ether)*

Data for **56**: (80%, white solid).  ^1^H-NMR (CDCl_3_, 400 MHz): δ = 8.03 (d, 1H), 7.7-7.6 (m, 2H), 7.17 (d, 1H), 6.98 (s, 2H), 6.75 (d, 4H), 5.87 (s, 4H), 4.11 (s, 4H), 1.24 (s, 18H) ppm.  ^13^C-NMR (CDCl_3_, 100 MHz): δ = 170.2, 169.2, 158.2, 152.9, 152.2, 135.1, 129.9, 129.4, 126.6, 125.2, 123.9, 113.4, 112.7, 103.6, 85.0, 82.4, 74.9, 60.4, 27.3 ppm. HRMS (APCI): calc’d for MH^+^ = C_34_H_36_O_11_H: 621.2330, found: 621.2327.

*Fluorescein bis*((*2-methoxy*)*pentanyloxymethyl ether)* (presumably racemic mixture)

Data for **57**: (79%, white solid).  ^1^H-NMR (CDCl_3_, 400 MHz): δ = 8.03 (d, 1H), 7.7-7.6 (m, 2H), 7.14 (d, 1H), 6.97  (m, 2H), 6.74 (s, 4H), 5.87 (dd, 4H), 3.81 (t, 2H), 3.37 (s, 6 H), 1.71 (m, 4H), 1.42-1.37 (m, 4H), 0.87 (t, 6H) ppm.  ^13^C-NMR (CDCl_3_, 100 MHz): δ = 171.8, 169.2, 158.2, 152.9, 152.2, 135.1, 129.9, 129.4, 126.6, 125.2, 123.8, 112.8, 112.7, 103.7, 85.1, 82.3, 80.2, 58.3, 34.7, 18.3, 13.7 ppm. HRMS (ESI): calc’d for MH^+^ C_34_H_36_O_11_Na: 643.2155; found: 643.2148.

*Fluorescein bis*((*2-isobutoxy)acetyloxymethyl ether)*

Data for **59**: (16%, white solid).  ^1^H-NMR (CDCl_3_, 400 MHz): δ = 8.06 (d, 1H), 7.72-7.64 (m, 2H), 7.17 (d, 1H), 6.98 (s, 2H), 6.76 (s, 4H), 5.88 (s, 4H), 4.16 (s, 4H), 3.31 (d, 4H), 1.97-1.87 (m, 2H), 0.93 (d, 12H) ppm.  ^13^C-NMR (CDCl_3_, 100 MHz): δ = 169.6, 169.1, 158.2, 152.9, 152.2, 135.1, 129.9, 129.4, 126.6, 125.2, 123.8, 113.5, 112.7, 103.6, 85.0, 82.3, 78.8, 68.2, 28.4, 19.2 ppm. HRMS (ESI): calc’d for MH^+^ C_34_H_36_O_11_H: 621.2336; found: 621.2321.

*Fluorescein bis*((*n-butoxy*)*acetyloxymethyl ether)*

Data for **61**: (40%, white solid).  ^1^H-NMR (CDCl_3_, 400 MHz): δ = 8.06 (d, 1H), 7.72-7.64 (m, 2H), 7.17 (d, 1H), 6.98 (s, 2H), 6.76 (s, 4H), 5.88 (s, 4H), 4.16 (s, 4H), 3.55 (t, 4H), 1.65-1.58 (m, 4H), 1.44-1.35 (m, 4H), 0.93 (t, 6H) ppm.  ^13^C-NMR (CDCl_3_, 100 MHz): δ = 169.6, 169.2, 158.2, 152.9, 152.2, 135.1, 129.9, 129.4, 126.6, 125.2, 123.8, 113.5, 112.7, 103.6, 85.0, 82.3, 71.9, 68.0, 31.5, 19.1, 13.8 ppm. HRMS (ESI): calc’d for MH^+^ C_34_H_36_O_11_H: 621.2336; found: 621.2309.

*Fluorescein bis((3-allyloxy)propanyloxymethyl ether)*

Data for **64**: (16%, white solid).  ^1^H-NMR (CDCl_3_, 400 MHz): δ = 8.05 (dm, 1H), 7.7-7.6 (m, 2H), 7.17 (dm, 1H), 6.98 (t, 2H), 6.75 (d, 4H), 5.9-5.8 (m, 6H), 5.3-5.1 (m, 4H), 3.98 (dt, *J* = 1.3, 5.5 Hz, 4H), 3.74 (t, *J =* 6.1 Hz, 4H), 2.69 (t, *J =* 6.1 Hz, 4H) ppm. ^13^C-NMR (CDCl_3_, 100 MHz): δ = 170.5, 169.2, 158.3, 152.9, 152.2, 135.1, 134.4, 129.9, 129.4, 126.6, 125.2, 123.9, 117.3, 113.3, 112.7, 103.6, 85.1, 82.4, 72.1, 65.01 35.1 ppm. HRMS (ESI): calc’d for MH^+^ C_34_H_32_O_11_H: 617.2017; found: 617.2018; calc’d for MNa^+^ C_34_H_32_O_11_Na: 639.1837; found: 639.1837.

*Fluorescein bis((3-isopropoxy)propanyloxymethyl ether)*

Data for **65**: (52%, white solid).  ^1^H-NMR (CDCl_3_, 400 MHz): δ = 8.05 (dm, 1H), 7.71-7.63 (m, 2H), 7.16 (dm, 1H), 6.99 (t, 2H), 6.75 (d, 4H), 5.83 (s, 4H), 3.72 (t, 4H), 3.58 (sep, 2H), 2.65 (t, 4H), 1.12 (d, 12H) ppm.  ^13^C-NMR (CDCl_3_, 100 MHz): δ = 170.7, 169.2, 158.4, 153.0, 152.2, 135.1, 129.9, 129.3, 126.6, 125.1, 123.8, 113.3, 112.7, 103.5, 85.0, 82.4, 71.9, 63.1, 35.5, 22.0 ppm. HRMS (ESI): calc’d for MNa^+^ C_34_H_36_O_11_Na: 643.2155; found: 643.2164.

*Fluorescein bis*((*3-isobutoxy*)*propanyloxymethyl ether)*

Data for **66**: (67%, white solid).  ^1^H-NMR (CDCl_3_, 400 MHz): δ = 8.05 (dm, 1H), 7.72-7.63 (m, 2H), 7.17 (d,m 1H), 6.99 (t, 2H), 6.75 (d, 4H), 5.83 (s 4H), 3.72 (t, 4H), 3.19 (d, 4H), 2.67 (t, 4H), 1.82 (sep, 2H), 0.86 (d, 12H) ppm.  ^13^C-NMR (CDCl_3_, 100 MHz): δ = 170.6, 169.2, 158.4, 153.0, 152.2, 135.1, 129.9, 129.3, 126.6, 125.1, 123.8, 113.3, 112.7, 103.5, 85.0, 82.4, 78.1, 65.8, 35.2, 28.3, 19.3 ppm. HRMS (ESI): calc’d for MNa^+^ C_36_H_40_O_11_Na: 671.2468; found: 671.2473.

*Fluorescein bis((4-ethoxy)butanyloxymethyl ether)*

Data for **68**: (57%, white solid).  ^1^H-NMR (CDCl_3_, 400 MHz): δ = 8.05 (dm, 1H), 7.72-7.64 (m, 2H), 7.18 (dm, 1H), 6.99 (t, 2H), 6.75 (d, 4H), 5.81 (s, 4H), 3.48-3.43 (m, 8H), 2.51 (t, 4H), 1.97-1.90 (m, 4H), 1.18 (t, 6H) ppm.  ^13^C-NMR (CDCl_3_, 100 MHz): δ = 172.3, 169.2, 158.4, 153.0, 152.2, 135.1, 129.9, 129.4, 126.6, 125.1, 123.9, 113.2, 112.7, 103.5, 84.9, 82.4, 69.1, 66.2, 31.1, 24.9, 15.1 ppm. HRMS (ESI): calc’d for MH^+^ C_34_H_36_O_11_H: 621.2336; found: 621.2335; calc’d for MNa^+^ C_34_H_36_O_11_Na: 643.2155; found: 643.2146.

*Fluorescein bis((4-isopropoxy)butanyloxymethyl ether)*

Data for **69**: (30%, white solid).  ^1^H-NMR (CDCl_3_, 400 MHz): δ = 8.05 (dm, 1H), 7.72-7.63 (m, 2H), 7.18 (d, 1H), 6.99 (t, 2H), 6.75 (d, 4H), 5.81 (s, 4H), 3.52 (sep, 2H), 3.44 (t, 4H), 2.51 (t, 4H), 1.95-1.88 (m, 4H), 1.12 (d, 12H) ppm. ^13^C-NMR (CDCl_3_, 100 MHz): δ = 172.4, 169.2, 158.4, 153.0, 152.2, 135.1, 129.9, 129.4, 126.6, 125.1, 123.9, 113.2, 112.7, 103.5, 84.8, 82.4, 71.5, 66.6, 31.1, 25.2, 22.0 ppm. HRMS (ESI): calc’d for MNa^+^ C_36_H_40_O_11_Na: 671.2468; found: 671.2319.

*Fluorescein bis*(*3-methylbut-2-enyloxymethyl ether)*

Data for **84**: (54%, white solid).  ^1^H-NMR (CDCl_3_, 400 MHz): δ = 8.04 (dm, 1H), 7.71-7.63 (m, 2H), 7.17 (dm, 1H), 7.00 (d, 2H), 6.78-6.73 (m, 4H), 5.84 (s, 4H), 5.75-5.74 (m, 2H), 2.22 (d, *J*=1.2Hz, 6H), 1.94 (d, *J*=1.3Hz, 6H) ppm. ^13^C-NMR (CDCl_3_, 100 MHz): δ = 169.3, 164.9, 160.3, 158.6, 153.0, 152.2, 135.1, 129.8, 129.3, 126.6, 125.1, 123.9, 114.9, 113.0, 112.7, 103.5, 84.3, 82.6, 27.6, 20.6 ppm. HRMS (ESI): calc’d for MH^+^ C_32_H_28_O_9_H: 557.1812; found: 557.1795.

*Fluorescein bis((2E)-2-methylbut-2-enyloxymethyl ether)*

Data for **85**: (79%, white solid).  ^1^H-NMR (CDCl_3_, 400 MHz): δ = 8.03 (d, 1H), 7.7-7.6 (m, 2H), 7.16 (d, 1H), 6.98-6.95 (m, 4H), 6.77-6.71 (m, 4H), 5.86 (s, 4H), 1.86 (s, 6H), 1.81 (d, 6H) ppm. ^13^C-NMR (CDCl_3_, 100 MHz): δ = 169.2, 166.5, 158.6, 152.9, 152.3, 139.6, 135.1, 129.9, 129.4, 127.8, 126.7, 125.1, 123.9, 113.1, 112.6, 103.5, 85.2, 82.6, 14.6, 12.0 ppm. HRMS: calc’d for MH^+^ = C_32_H_28_O_9_H: 557.1806, found: 557.1808.

*Fluorescein bis(2,3-dimethylbut-2-enyloxymethyl ether)*

Data for **86**: (66%, white solid).  ^1^H-NMR (CDCl_3_, 400 MHz): δ = 8.05 (d, 1H), 7.7-7.6 (m, 2H), 7.18 (d, 1H), 7.00 (d, 2H), 6.79-6.73 (m, 4H), 5.87 (s, 4H), 2.08 (d, 6H), 1.89 (s, 6H), 1.87 (s, 6H) ppm.  ^13^C-NMR (CDCl_3_, 100 MHz): δ = 169.3, 167.5, 158.6, 153.0, 152.3, 147.8, 135.1, 129.9, 129.3, 126.7, 125.1, 123.9, 121.2, 113.1, 112.7, 103.6, 84.9, 82.6, 23.2, 23.1, 15.5 ppm. HRMS: calc’d for MH^+^ = C_34_H_32_O_9_H: 585.2119, found: 585.2118; calc’d for MNa^+^ = C_34_H_32_O_9_S_2_Na: 607.1939, found: 607.1938.

*Fluorescein bis((2E)-3-chloroacryloxymethyl ether)*

Data for **87**: (18%, white solid). ^1^H-NMR (CDCl_3_, 400 MHz): δ = 8.04 (d, 1H), 7.7-7.6 (m, 2H), 7.47 (d, *J*=13.5Hz, 2H), 7.16 (d, 1H), 6.96 (d, 2H), 6.74 (m, 4H), 6.29 (d, *J*=13.5Hz, 2H), 5.87 (s, 4H) ppm. ^13^C-NMR (CDCl_3_, 100 MHz): δ = 169.2, 162.6, 158.2, 152.9, 152.2, 139.9, 135.2, 129.9, 129.5, 126.6, 125.2, 123.9, 123.8, 113.5, 112.7, 103.5, 85.2, 82.3 ppm. HRMS (ESI): calc’d for MH^+^ C_28_H_18_O_9_Cl_2_H: 569.0400; found: 569.0401; calc’d for MNa^+^ C_28_H_18_O_9_Cl_2_Na: 591.0220; found: 591.0220.

*Fluorescein b bis((2Z)-3-chloroacryloxymethyl ether)*

Data for **88**: (29%, white solid).  ^1^H-NMR (CDCl_3_, 400 MHz): δ = 8.03 (d, 1H), 7.7-7.6 (m, 2H), 7.16 (d, 1H), 6.99 (m, 2H), 6.83 (d, *J* = 8.2 Hz, 2H), 6.75 (m, 4H), 6.25 (d, *J* = 8.2 Hz, 2H), 5.89 (s, 4H) ppm.  ^13^C-NMR (CDCl_3_, 100 MHz): δ = 169.2, 161.9, 158.3, 152.9, 152.2, 135.2, 135.1, 129.9, 129.4, 126.6, 125.2, 123.9, 120.3, 113.4, 112.7, 103.6, 85.1, 82.3 ppm. HRMS (ESI): calc’d for MH^+^ C_28_H_18_O_9_Cl_2_H: 569.0401; found: 569.0402; calc’d for MNa^+^ C_28_H_18_O_9_Cl_2_Na: 591.0220; found: 591.0220.

*Fluorescein bis((E)-3-acetylacryloxymethyl ether)*

Data for **89**: (98%, white solid).  ^1^H-NMR (CDCl_3_, 400 MHz): δ = 8.05 (d, 1H), 7.73-7.64 (m, 2H), 7.18 (d, 1H), 7.11(d, *J*=16Hz, 2H), 7.00 (s, 2H), 6.78 (s, 4H), 6.69 (d, *J*=16Hz, 2H), 5.94 (s, 4H), 2.39 (s, 6H) ppm.  ^13^C-NMR (CDCl_3_, 100 MHz): δ = 197.2, 169.2, 164.2, 158.1, 152.8, 152.2, 141.4, 135.2, 130.1, 130.0, 129.5, 126.5, 125.2, 123.9, 113.6, 112.7, 103.6, 85.6, 82.2, 28.3 ppm. HRMS (ESI): calc’d for MH^+^ C_32_H_24_O_11_H: 585.1397; found: 585.1411.

*Fluorescein bis((2E)-hex-2-enyloxymethyl ether)*

Data for **91**: (54%, white solid).  ^1^H-NMR (CDCl_3_, 400 MHz): δ = 8.04 (dm, 1H), 7.7-7.6 (m, 2H), 7.17 (d, 1H), 7.10 (dt, *J*’s = 16.4 & 7.0 Hz, 2H), 7.00 (dm, 2H), 6.79-6.74 (m, 4H), 5.87 (dt, *J*’s = 15.7 & 1.6 Hz, 2H), 5.87 (s, 4H), 2.22 (qd, *J*’s = 7.2 & 1.4 Hz, 4H), 1.56-1.47 (m, 4H), 0.95 (t, 6H) ppm.  ^13^C-NMR (CDCl_3_, 100 MHz): δ = 169.2, 165.1, 158.5, 152.9, 152.2, 152.0, 135.1, 129.9, 129.3, 126.6, 125.1, 123.9, 120.2, 113.2, 112.7, 103.5, 84.9, 82.5, 34.4, 21.1, 13.7 ppm. HRMS: calc’d for MH^+^ = C_34_H_32_O_9_H: 585.2119, found: 585.2119; calc’d for MNa^+^ = C_34_H_32_O_9_Na: 607.1939, found: 607.1940.

*Fluorescein bis*(*oleyloxymethyl ether)*

Data for **92**: (83%, white solid).  ^1^H-NMR (CDCl_3_, 400 MHz): δ = 8.03 (d, 1H), 7.7-7.6 (m, 2H), 7.15 (d, 1H), 6.96 (s, 2H), 6.73 (m, 4H), 5.78 (s, 4H), 5.37-5.28 (m, 4H), 2.37 (t, 4H), 2.05-1.95 (m, 8H), 1.65-1.62 (m, 4H), 1.4-1.2 (m, 40H), 0.87 (t, 6H) ppm.  ^13^C-NMR (CDCl_3_, 100 MHz): δ = 172.6, 169.2, 158.4, 153.0, 152.2, 135.1, 130.0, 129.9, 129.7, 129.4, 126.7, 125.2, 123.9, 113.2, 112.7, 103.5, 84.8, 82.4, 36.6, 34.1, 31.9, 29.8, 29.7, 29.5, 29.3, 29.1, 29.06, 29.0, 27.2, 27.1, 24.7, 24.6, 23.3, 22.7, 14.1 ppm. HRMS (APCI): calc’d for MH^+^ = C_58_H_80_O_9_H: 921.5875, found: 921.5874.

*Fluorescein bis(propynyloxymethyl ether)*

Data for **93**: (74%, white solid).  ^1^H-NMR (CDCl_3_, 400 MHz): δ = 8.03 (d, 1H), 7.7-7.6 (m, 2H), 7.15 (d, 1H), 6.98 (m, 2H), 6.76 (m, 4H), 5.86 (s, 4H), 3.00 (s, 2H) ppm.  ^13^C-NMR (CDCl_3_, 100 MHz): δ = 169.2, 158.0, 152.9, 152.2, 151.3, 135.2, 130.0, 129.5, 126.5, 125.2, 123.9, 113.8, 112.8, 103.8, 86.0, 82.2, 77.2, 73.8 ppm. HRMS: calc’d for MH^+^ = C_28_H_16_O_9_H: 497.0867, found: 497.0868; calc’d for MNa^+^ = C_28_H_16_O_9_Na: 519.0687, found: 519.0688.

*Fluorescein bis((1-pent-2-ynyl)oxymethyl ether)*

Data for **95**: (40%, white solid).  ^1^H-NMR (CDCl_3_, 400 MHz): δ = 8.03 (d, 1H), 7.7-7.6 (m, 2H), 7.15 (d, 1H), 6.98 (s, 2H), 6.75 (s, 4H), 5.83 (s, 4H), 2.36 (q, 4H), 1.21 (t, 6H) ppm.  ^13^C-NMR (CDCl_3_, 100 MHz): δ = 169.2, 158.2, 153.0, 152.3, 152.2, 135.1, 129.9, 129.5, 126.6, 125.2, 123.9, 113.5, 112.7, 103.7, 93.2, 85.6, 82.3, 71.7, 12.52, 12.36 ppm. HRMS (APCI): calc’d for MH^+^ = C_32_H_24_O_9_H: 553.1493, found: 553.1492; calc’d for MNa^+^ = C_32_H_24_O_9_Na: 575.1313, found: 575.1311.

*Fluorescein bis(4-acetylbutanyloxymethyl ether)*

Data for **96**: (37%, white solid). ^1^H-NMR (CDCl_3_, 400 MHz): δ = 8.05 (d, 1H), 7.7-7.6 (m, 2H), 7.18 (d, 1H), 6.98 (s, 2H), 6.76 (s, 4H), 5.81 (s, 4H), 2.52 (t, 4H), 2.45 (t, 4H), 2.12 (s, 6H), 1.96-1.89 (m, 4H) ppm. ^13^C-NMR (CDCl_3_, 100 MHz): δ = 207.7, 171.9, 169.2, 158.3, 152.9, 152.2, 135.1, 129.9, 129.4, 126.6, 125.2, 123.9, 113.3, 112.7, 103.5, 84.8, 82.4, 42.1, 33.0, 29.9, 18.5 ppm. HRMS: calc’d for MH^+^ = C_34_H_36_O_11_H: 617.2023, found: 617.2007.

**Generation of *P. falciparum* and erythrocyte lysate**

Unsynchronized *P. falciparum* cultures were lysed with 1% saponin and washed in Dulbecco’s phosphate-buffered saline (dPBS) (Gibco) followed by lysis buffer containing 100 mM Tris-HCl (pH 7.5), 100 mM NaCl, 1 mM MgCl2, 1 mM DTT, 10% glycerol, and cOmplete EDTA-Free protease inhibitor cocktail (Roche). Purified *P. falciparum* pellets were resuspended in lysis buffer and lysed by sonication (Fisherbrand™ Model 120 Sonic Dismembrator; 3 cycles, 10 second pulse, 10 second rest, 40% power). Cell debris were removed by centrifugation and the remaining *P. falciparum* lysate was used in downstream assays. Erythrocyte lysate was generated by the same method, excluding the saponin lysis step.

**Peptide mass spetrometry**

*Tryptic digestion*

Excised gel bands were cut into 1 mm^3^ cubes, destained with 50% Methanol/1.25% Acetic Acid, reduced with 5 mM DTT (Dithiothreitol) (Thermo), and alkylated with 20 mM iodoacetamide (Sigma). Gel pieces were then washed with 20 mM ammonium bicarbonate(Sigma) and dehydrated with acetonitrile (Fisher). Trypsin (Promega)(5ng/uL in 20 mM ammonium bicarbonate) was added to the gel pieces and proteolysis was allowed to proceed overnight at 37 ºC. Peptides were extracted with 0.3% triflouroacetic acid (J.T.Baker), followed by 50% acetonitrile. Extracts were combined and the volume was reduced by vacuum centrifugation. The resulting peptides were de-salted, dried by vacuum centrifugation and reconstituted in 0.1% TFA containing iRT peptides (Biognosys Schlieren, Switzerland).

*Mass Spectrometry Data Acquisition*

Samples were analyzed on a Q-Exactive HF mass spectrometer (Thermofisher Scientific San Jose, CA) coupled with an Ultimate 3000 nano UPLC system and an EasySpray source. Peptides were loaded onto an Acclaim PepMap 100 75um x 2cm trap column (Thermo) at 5uL/min and separated by reverse phase (RP)-HPLC on a nanocapillary column, 75 μm id × 50cm 2um PepMap RSLC C18 column (Thermo). Mobile phase A consisted of 0.1% formic acid and mobile phase B of 0.1% formic acid/acetonitrile. Peptides were eluted into the mass spectrometer at 300 nL/min with each RP-LC run comprising a 90-minute gradient from 3% B to 45% B.

The mass spectrometer was set to repetitively scan m/z from 300 to 1400 (R = 240,000) followed by data-dependent MS/MS scans on the twenty most abundant ions, minimum AGC 1e4, dynamic exclusion with a repeat count of 1, repeat duration of 30s, and resolution of 15000. The AGC target value was 3e6 and 1e5, for full and MSn scans, respectively. MSn injection time was 160 ms. Rejection of unassigned and 1+,6-8 charge states was set.

*System suitability and quality control*

The suitability of Q Exactive HF instrument was monitored using QuiC software (Biognosys, Schlieren, Switzerland) for the analysis of the spiked-in iRT peptides. Meanwhile, as a measure for quality control, we injected standard E. coli protein digest prior to and after injecting sample set, and collected the data in the Data Dependent Acquisition (DDA) mode. The collected DDA data were analyzed in MaxQuant^8^ and the output was subsequently visualized using the PTXQC^9^ package to track the quality of the instrumentation.

*Database Searching*

All MS/MS samples were analyzed using MSFragger (The Nesvizhskii Lab, 1301 Catherine, 4237 Medical Science I, Ann Arbor, MI 48109). MSFragger was set up to search a reverse concatenated Pfalciparum_211123 database (5385 entries) assuming the digestion enzyme trypsin. MSFragger was searched with a fragment ion mass tolerance of 20 PPM and a parent ion tolerance of 20 PPM. Carbamidomethyl of cysteine was specified in MSFragger as a fixed modification. Oxidation of methionine was specified in MSFragger as a variable modification.

Scaffold (version Scaffold_5.1.0, Proteome Software Inc., Portland, OR) was used to validate MS/MS based peptide and protein identifications. Peptide identifications were accepted if they could be established at greater than 95.0% probability by the Percolator posterior error probability calculation^10^. Protein identifications were accepted if they could be established at greater than 99.0% probability and contained at least 1 identified peptide. Protein probabilities were assigned by the Protein Prophet algorithm^11^. Proteins that contained similar peptides and could not be differentiated based on MS/MS analysis alone were grouped to satisfy the principles of parsimony. Proteins sharing significant peptide evidence were grouped into clusters.

**Expression of recombinant APEH**

A 6x histidine tagged full length cDNA sequence of APEH (SinoBiological) was expressed in 293F cells using the pcDNA3.1 vector. The final plasmid sequence was confirmed by nanopore sequencing (Plasmidsaurus).  Cells were harvested after 48 hours of expression, pelleted, and lysed by sonication. rAPEH was purified using TALON® Metal Affinity Resin (Takara) in phosphate buffered saline (PBS) with an additional 100 mM NaCl, 2 mM MgCl2, 2 mM KCl and 10% glycerol. Complete Edta-Free protease inhibitor cocktail (Roche) and benzonase (Sigma) were added to the lysis buffer. Lysis and washes were performed with 5 mM imidazole and elution with 150 mM imidazole. Protein purity was confirmed by SDS-PAGE.

**Activation of POM-ERJ measured by mass spectrometry**

Reactions containing dPBS with 2 mM MgCl2, 10 µM POM-ERJ, and either 15 ng/µl rAPEH, 15 ng/µl rAPEH and 400 nM AA74-1, or buffer were incubated at 37 °C and sampled at 0, 1, 2, 4, and 6 hours. At each timepoint, 25 µl of reaction was removed, quenched with 100 µl of cold acetonitrile, and stored at -20 °C.

Quenched reaction mixtures were analyzed as previously described ^12^with the following modifications.  To the quenched samples, 10 µl of 200 ng/mL enalapril in methanol was added as the internal standard and then analyzed on a Sciex 6500 Q  LC-MSMS system. Standards were evaluated over a range of 500 to 10,000 ng/mL . The MRM transitions for enalapril and POM-ERJ were *m/z* 377 > 91 and 424 > 364, respectively. A Phenomenex Kinetex polar C18 column (50 x 2.1 mm, 2.6 μm) was used for chromatographic separation. Mobile phases were 0.1% formic acid in water and acetonitrile with a flow rate of 0.4 mL/min. Peak areas were integrated using Analyst Software (Sciex, Framingham, MA).

**APEH activity in intact erythrocytes**

Erythrocytes at 2% hematocrit in culture media were incubated for 24 hours with AA74-1 at concentrations of 500 nM to 10 µM. For washed samples, cells were washed three times in culture media. Samples were taken at 0, 24, 48, and 72 hours, and APEH activity was quantified by measuring rates of AANA hydrolysis (22). For each timepoint, 50 µl of cells were removed and washed three times with dPBS + 1.25 mM MgCl_2_. Cells were then mixed with dPBS with 1.25 mM MgCl_2_ and 5 mM acetylalanine 4-nitroanilide (AANA) (Carbosynth). The rate of hydrolysis of AANA was monitored by measuring the increase in absorbance at 410 nm. Percent APEH activity was determined by comparing the rate of hydrolysis to that of untreated erythrocytes.

***P. falciparum* growth inhibition by AA74-1 with or without washout**

Uninfected erythrocytes in culture media were incubated with AA74-1 at concentrations from 9.8 nM to 5 µM for 24 hours in 96 well plates. For washout, erythrocytes were then washed three times and resuspended in culture media without AA74-1. Magnetically isolated *P. falciparum* schizonts were added to pretreated erythrocytes at 1% parasitemia. Cultures were incubated for 72 hours days and parasite growth quantified by measuring DNA content using PicoGreen (Life Technologies) as previously described (7). EC_50_ values were calculated by nonlinear regression analysis using GraphPad Prism software.
